# Transcription factors ERα and Sox2 have differing multiphasic DNA and RNA binding mechanisms

**DOI:** 10.1101/2024.03.18.585577

**Authors:** Wayne O. Hemphill, Halley R. Steiner, Jackson R. Kominsky, Deborah S. Wuttke, Thomas R. Cech

## Abstract

Many transcription factors (TFs) have been shown to bind RNA, leading to open questions regarding the mechanism(s) of this RNA binding and its role in regulating TF activities. Here we use biophysical assays to interrogate the k_on_, k_off_, and K_d_ for DNA and RNA binding of two model human transcription factors, ERα and Sox2. Unexpectedly, we found that both proteins exhibited multiphasic nucleic acid binding kinetics. We propose that Sox2 RNA and DNA multiphasic binding kinetics could be explained by a conventional model for sequential Sox2 monomer association and dissociation. In contrast, ERα nucleic acid binding exhibited biphasic dissociation paired with novel triphasic association behavior, where two apparent binding transitions are separated by a 10-20 min “lag” phase depending on protein concentration. We considered several conventional models for the observed kinetic behavior, none of which adequately explained all the ERα nucleic acid binding data. Instead, simulations with a model incorporating sequential ERα monomer association, ERα nucleic acid complex isomerization, and product “feedback” on isomerization rate recapitulated the general kinetic trends for both ERα DNA and RNA binding. Collectively, our findings reveal that Sox2 and ERα bind RNA and DNA with previously unappreciated multiphasic binding kinetics, and that their reaction mechanisms differ with ERα binding nucleic acids via a novel reaction mechanism.

## INTRODUCTION

The human genome encodes ∼1500 transcription factors (TFs) (Ignatieva et al. 2015; Zhang et al. 2012; Wingender et al. 2013, 2015), which direct cell-type specificity and gene expression programs by interacting with a multitude of binding partners (Spitz and Furlong 2012). TFs modulate transcription by utilizing their DNA-binding domains to stably interact with DNA elements, such as promoters and enhancers, with sequence specificity, and subsequently recruit various coactivator and repressor proteins via their effector domains (Schwabe et al. 1993; Frietze and Farnham 2011). However, the site of transcription is immersed in more than DNA and protein – it’s also crowded with RNA. Thousands of RNA species are produced at loci where TFs are bound, such as mRNA, enhancer RNAs, promoter antisense RNAs, and chromatin-enriched long noncoding RNAs (Han and Li 2022; Yang et al. 2021; Werner and Ruthenburg 2015). Additionally, long noncoding RNAs transcribed distally, even kilobases away, are capable of engaging in long-range interactions with chromatin (Mishra and Kanduri 2019; Rinn and Chang 2020). The prevalence of RNA at chromatin begs the question of whether RNA plays a direct role in regulating TFs.

Many TFs have been shown to bind RNA (G Hendrickson et al. 2016; Khalil et al. 2009; Skalska et al. 2021; Oksuz et al. 2023; Hudson and Ortlund 2014; Parsonnet et al. 2019), raising the question of whether RNA directly regulates TF function. In many cases, the TF’s RNA binding domains are adjacent to their DNA binding domains (Oksuz et al. 2023). Estrogen receptor α (ERα) (Steiner et al. 2022) and sex determining region Y box 2 (Sox2) (Holmes et al. 2020) are two such TFs that bind DNA and RNA competitively with tight affinities, suggesting potential biological relevance for the RNA binding activity. ERα and Sox2 can therefore be used as model systems to study RNA regulation of TF activities. Although RNA-DNA competition experiments provide some useful information, a detailed investigation of the mechanism(s) and kinetics for TF polynucleotide association and dissociation is critical for understanding how TFs could be regulated by RNA binding.

ERα is a ligand-activated TF, which functions as the nuclear receptor for estrogen, a hormone that dictates reproductive development (Björnström and Sjöberg 2005; Deroo and Korach 2006) (mouse studies reviewed in ref (Hewitt and Korach 2018)). Abnormal ERα signaling leads to a variety of diseases such as metabolic and cardiovascular disease, neurodegeneration, and inflammation (Jia et al. 2015). Additionally, ERα is aberrantly expressed in 80% of breast cancers, making it a recurrent therapeutic target (Alluri et al. 2014). ERα is a 595 amino acid polypeptide (∼66 kDa) comprised of six domains, including DNA-binding, ligand-binding and transcriptional activation domains (Hewitt and Korach 2018; Ponglikitmongkol et al. 1988). Its DNA-binding domain (DBD) facilitates sequence-specific DNA binding to the palindromic estrogen response element (ERE) motif (GGTCAnnnTGACC) and binds as a dimer via its two zinc finger elements (Kuntz and Shapiro 1997; Helsen et al. 2012; Schwabe et al. 1993). The hinge region sits just C-terminal of the DBD, and recent work has demonstrated that part of the hinge region is critical for RNA binding (but dispensable for DNA binding). ERα uses a combination of the DBD and hinge elements to preferentially bind hairpin RNA (hRNA) with no apparent sequence specificity (Steiner et al. 2022; Xu et al. 2021). While *in vitro* experiments indicate that ERα RNA and DNA binding are competitive, and ERα has been shown to interact with RNA *in vivo* (Xu et al. 2021; Nassa et al. 2019), the question of how RNA may regulate ERα-DNA interactions *in vivo* remains an active area of investigation (Steiner et al. 2022).

Sox2, a member of the SoxB1 TF family, regulates pluripotency in embryonic stem cells via expression of the pluripotency-associated TFs Oct4 and Nanog and via repression of lineage-specific genes (Avilion et al. 2003; Zhang and Cui 2014; Chew et al. 2005). Additionally, Sox2 is critical for differentiating pluripotent stem cells to neural progenitors and maintaining the properties of neural progenitor stem cells (Zhang and Cui 2014). In mice, deletion of Sox2 is embryonic lethal (Avilion et al. 2003), while knockout in adult mice leads to the loss of hippocampal neurogenesis (Favaro et al. 2009). In humans, mutations in Sox2 have been associated with eye defects such as bilateral anophthalmia and microphthalmia (Fantes et al. 2003; Chassaing et al. 2014), as well as cognitive abnormalities (Sisodiya et al. 2006; Kelberman et al. 2006). Functional Sox2 contains 317 amino acids partitioned into two key domains (Nowling et al. 2000). The Sox2 high mobility group (HMG) domain binds DNA at the minor groove, and recognizes a species-specific sequence centered around four highly conserved nucleotides (CCCA<u>TTGT</u>TC in humans) (Grosschedl et al. 1994; Dodonova et al. 2020; Schaefer and Lengerke 2020; Yesudhas et al. 2017; Hou et al. 2017; Weiss 2001). *In vivo* studies have suggested that lncRNAs interact directly with Sox2 to regulate its function(s) in stem cell pluripotency (Ng et al. 2012). Subsequent *in vitro* findings show that the Sox2 HMG domain preferentially binds the double-stranded RNA within hRNA with no apparent sequence specificity (Holmes et al. 2020). Like ERα, Sox2 HMG domain binding to RNA and DNA was found to be competitive (Holmes et al. 2020). However, another study suggests that a novel RNA-binding module C-terminal of the HMG domain also contributes to Sox2 RNA binding, and that Sox2 can stably bind RNA and DNA simultaneously (Hou et al. 2020).

To determine the mechanisms of RNA and DNA binding on the Sox2 and ERα binding surfaces, we used fluorescence polarization (FP) and surface plasmon resonance (SPR) to measure their RNA and DNA association and dissociation kinetics. In contrast to the expectation for a simple binding reaction, both TFs exhibited complex multiphasic association and dissociation kinetics from RNA and DNA. We evaluated several common models for multiphasic association and/or dissociation to describe the observed kinetics for the two TF interactions with RNA and DNA. These findings reveal a previously unappreciated level of complexity in the ERα and Sox2 interactions with nucleic acids, and they suggest that the two TFs achieve multiphasic kinetics through different mechanisms.

## RESULTS

### ERα_DBD-Ext_ and Sox2_HMG_ equilibrium ligand binding

ERα_180-280_, a region of the protein containing the canonical DBD and a set of basic residues from the hinge region (ERα_DBD-Ext_), and recombinant Sox2_40-123_, the region of the protein containing the high mobility group (HMG) domain (Sox2_HMG_), were expressed and purified as previously described (Steiner et al. 2022; Holmes et al. 2020). We performed FP-based binding experiments with ERα_DBD-Ext_ and Sox2_HMG_ and a variety of dsDNA and RNA ligands to assess the binding affinities (Supplemental Figure 1) at the same experimental conditions used to measure binding kinetics.

For ERα_DBD-Ext_, we tested an 18-bp dsDNA containing its palindromic ERE recognition sequence (ERE dsDNA), a 15-bp dsDNA containing only half of its palindromic recognition sequence (ΔERE dsDNA), and a 37-nt hairpin RNA (hRNA) derived from the X-box binding protein 1 (XBP1) mRNA sequence (XBP1 hRNA) (Steiner et al. 2022). We found that ERα_DBD-Ext_ bound ERE with high affinity (K_d_^app^ ≈ 11 nM, see Table 1 for error) and positive cooperativity (n ≈ 2.1) (Supplemental Figure 1a and Table 1), while ERα_DBD-Ext_ bound ΔERE with comparable to higher affinity (K_d_^app^ ≈ 2.8 nM) and less to no positive cooperativity (n ≈ 1.5) (Supplemental Figure 1a and Table 1). ERα_DBD-Ext_ bound the XBP1 hRNA with lower affinity (K_d_^app^ ≈ 370 nM) and no apparent cooperativity (n ≈ 0.91) (Supplemental Figure 1a and Table 1). These findings are consistent with prior studies (Steiner et al. 2022). We note that the anisotropy dynamic range was less for the ΔERE versus ERE binding curve, consistent with a lower TF-DNA binding stoichiometry for ΔERE versus ERE, as expected from prior studies (Steiner et al. 2022; Schwabe et al. 1993).

**Table 1:** Kinetic Constant Values from FP-based ERα _DBD-Ext_ and Sox2_HMG_ Binding Experiments. Table includes apparent equilibrium dissociation constants (K_d_^app^) and Hill coefficients (n) from Supplemental Figure 1 (fit with Eq. 1.1-2), apparent initial association rate constants (k_on_^app^) from Figure 3, and apparent dissociation rate constants (k_off_^app^) from Figure 1 (fit with Eq. 3). Values are mean ± ½ range (across 2-3 independent experiments). Values without error are based on a single experiment. For K_d_^app^, values in parentheses are the percent signal contributions of the first transitions in two-transition binding regression, and for k_off_^app^, values in parentheses are the percent contributions of the fast or slow components in bi-exponential regression. Percentages are the averages across 2-3 independent experiments, or values from a single experiment if the associated rate constants have no error.

| Protein | Ligand | $K_d^{app}$ (nM) | n (Hill) | $k_{on}^{app}$ ($M^{-1}s^{-1}$) | $k_{off}^{app_{fast}}$ ($\times 10^{-2} s^{-1}$) | $k_{off}^{app_{slow}}$ ($\times 10^{-4} s^{-1}$) |
| --- | --- | --- | --- | --- | --- | --- |
| *ERα*<sub>DBD-Ext</sub> | ERE dsDNA | 11 ± 2.1 | 2.1 ± 0.30 | 5.8 ± 3.1 $\times 10^4$ | 4.7 (71%) | 7.3 (29%) |
| | ΔERE dsDNA | <sup>¥</sup>2.8 ± 0.6 | <sup>¥</sup>1.5 ± 0.22 | 8.7 ± 4.6 $\times 10^4$ | 8.9 ± 4.4 (60%) | 5.9 ± 1.1 (40%) |
| | XBP1 hRNA | 370 | 0.91 | 14 $\times 10^4$ | 5.6 ± 2.7 (64%) | 6.6 ± 1.2 (36%) |
| *Sox2*<sub>HMG</sub> | CBS dsDNA | <sup>†</sup>≤1 (54%), 420 ± 120 | <sup>†</sup> n/a | 0.75 $\times 10^6$ | 5.0 ± 2.3 (67%) | 13 ± 2.1 (33%) |
| | G4 RNA | <sup>†</sup>≤1 (44%), 110 ± 32 | <sup>†</sup> n/a | 1.1 $\times 10^6$ | 5.4 (73%) | 9.6 (27%) |
| | NBS dsDNA | 29 ± 9.1 | 0.57 ± 0.06 | n.d. | n.d. | n.d. |
| | hRNA | 53 ± 16 | 1.1 ± 0.21 | n.d. | n.d. | n.d. |
| | poly-A RNA | 40 ± 13 | 0.68 ± 0.11 | n.d. | n.d. | n.d. |

n/a = not applicable
n.d. = not determined
<sup>¥</sup> $K_d^{app}$ is close to ligand concentration; it's possible that $K_d < K_d^{app}$ and n (Hill coefficient) is artificially inflated
<sup>†</sup> Binding curves were fit with a two-transition equation, not Hill equation

For Sox2_HMG_, we tested a 10-bp dsDNA containing its cognate binding sequence (CBS dsDNA) (Holmes et al. 2020), and for comparison we also measured the binding affinities for a 19-bp dsDNA with a nonspecific binding sequence (NBS dsDNA), a 43-nt hairpin RNA (hRNA), a (G_3_A_2_)_4_ RNA that adopts a G-quadruplex (G4) structure (rG4), and a 20-nt poly-A RNA (rA20) (Supplemental Figure 1b and Table 1). Our findings indicate that Sox2_HMG_ binding to CBS dsDNA and G4 RNA were best described by a two-transition binding curve, while Sox2_HMG_ binding to NBS dsDNA, hRNA, and poly-A RNA fit well to a standard Hill binding equation. The Sox2_HMG_ CBS and rG4 high-affinity binding transitions both had K_d_^app^ ≤ 1 nM, being limited by the ligand concentration in our assays, while their lower-affinity binding transitions had K_d_^app^ of 420 and 110 nM, respectively. For CBS dsDNA, this was previously attributed to Sox2_HMG_ initial sequence-specific binding versus subsequent nonspecific binding (Holmes et al. 2020; Hamilton et al. 2022). Relative to the high-affinity binding transition, Sox2_HMG_ exhibited ≥30x greater affinity for CBS dsDNA and G4 RNA than for NBS dsDNA, hRNA, and poly(A) RNA (K_d_^app^ ≈ 29-53 nM). We also note that the Sox2_HMG_ NBS dsDNA and poly(A) RNA binding curves exhibited modest negative cooperativity (n ≈ 0.57-0.68), while Sox2_HMG_ bound the hRNA without apparent cooperativity (Supplemental Figure 1b and Table 1). All these findings are in agreement with prior studies (Hamilton et al. 2022; Holmes et al. 2020), validating the reagents and methods for the subsequent measurements below.

### ERα_DBD-Ext_ and Sox2_HMG_ ligand dissociation are multiphasic

We measured the ERα_DBD-Ext_ and Sox2_HMG_ ligand dissociation kinetics using FP-based competitive dissociation (FPCD) experiments. These involve pre-incubation of protein and fluorescently labeled nucleic acid followed by self-competition with unlabeled nucleic acid and observation of binding states by FP (Figure 1a). Contrary to the expectation for a simple binding scheme (i.e., protein + ligand ←→ protein-ligand), both ERα_DBD-Ext_ and Sox2_HMG_ exhibited multiphasic dissociation from all ligands that we tested (Figure 1b-c). For ERα_DBD-Ext_, the ligand dissociation curves were well fit by bi-exponential regression and produced similar rate constants for all the nucleic acids tested. The rate constants and other relevant parameter values for these regressions are summarized in Table 1.

**Figure 1:**
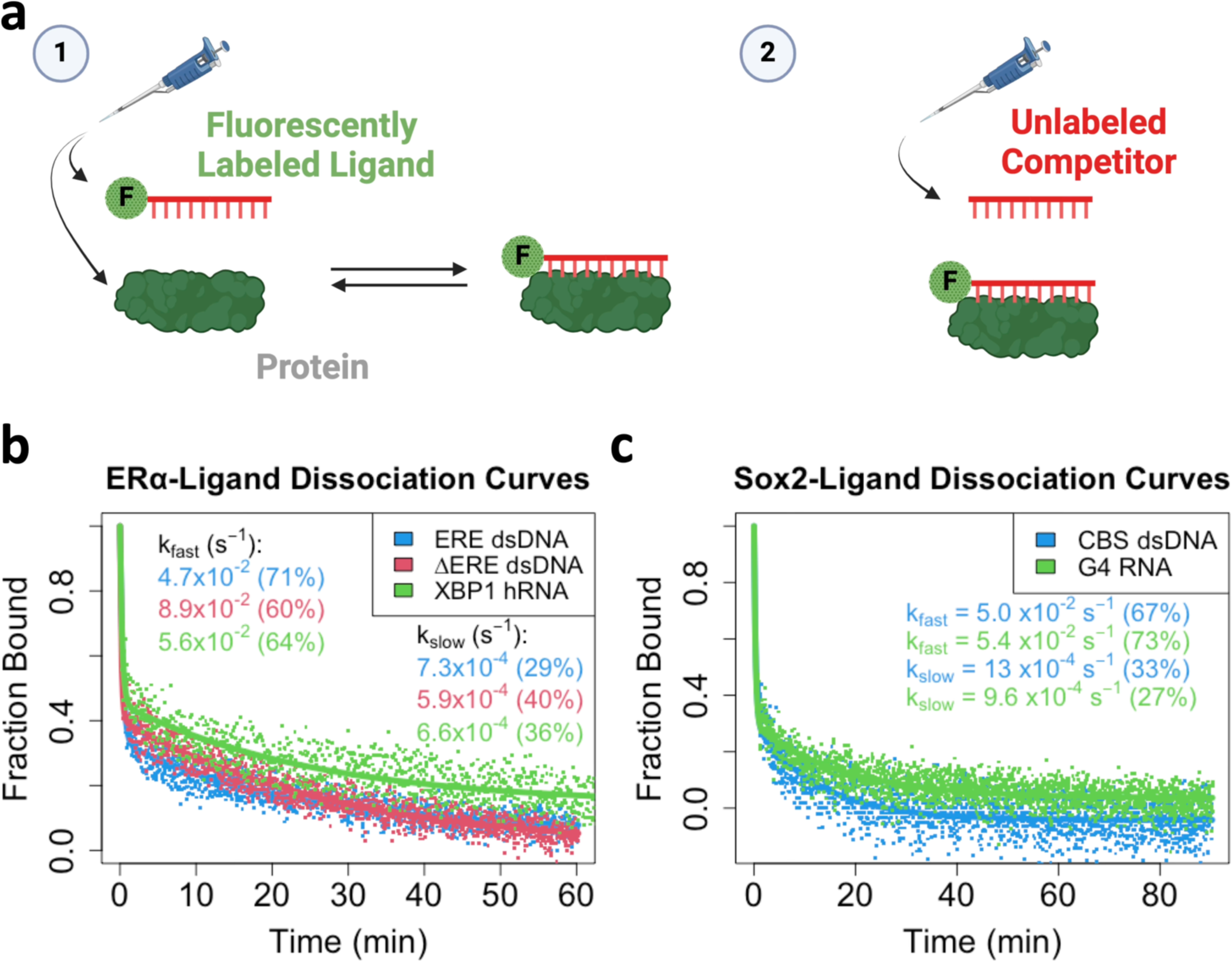
ERα_DBD-Ext_ and Sox2_HMG_ Exhibit Multiphasic Ligand Dissociation. **[a]** *Graphical summary of Fluorescence Polarization-based Competitive Dissociation (FPCD) experiments.* (1) Fluorescently labeled polynucleotide is mixed with protein and incubated at 4°C for a variable amount of time, then (2) an excess of unlabeled polynucleotide (i.e., competitor) is added to the protein-ligand reaction and polarization is monitored over time (at 4°C) to observe protein-ligand dissociation kinetics. **[b-c]** *Dissociation curves from FPCD experiments.* FPCD experiments (panel a) were performed using 5 nM ligand, 10 µM competitor, and 100-500 nM protein; protein-nucleic acid binding reactions were incubated long enough to reach equilibrium before competitor addition. Anisotropy traces were normalized to the internal controls to give ‘Fraction Bound’ over time, then normalized data fit with bi-exponential regression (Eq. 3) to determine rate constants. Dots are data points and solid lines are regression fits from a single experiment for each ligand. Rate constants and (in parenthesis) the percent contributions of fast versus slow components to the bi-exponential regression are reported with error in Table 1.

We then asked what could be producing the biphasic dissociation curves for our TF nucleic acid interactions. Previously, we demonstrated that direct transfer is used by multiple nucleic acid binding proteins to transfer between polynucleotide species through unstable ternary intermediates (Hemphill et al. 2023). Furthermore, at the competitor concentrations used in our FPCD experiments (Figure 1), protein-polynucleotide dissociation might occur via both direct transfer and intrinsic dissociation in comparable proportions (Hemphill et al. 2023). If the fast components of the ERα_DBD-Ext_ biphasic dissociation curves were the result of ligand displacement via direct transfer, their dissociation curves should become monophasic slow in the absence of competitor. To test this hypothesis, we induced complex dissociation by dilution rather than competitor addition with FP-based jump dilution (FPJD) experiments (Figure 2).

**Figure 2:**
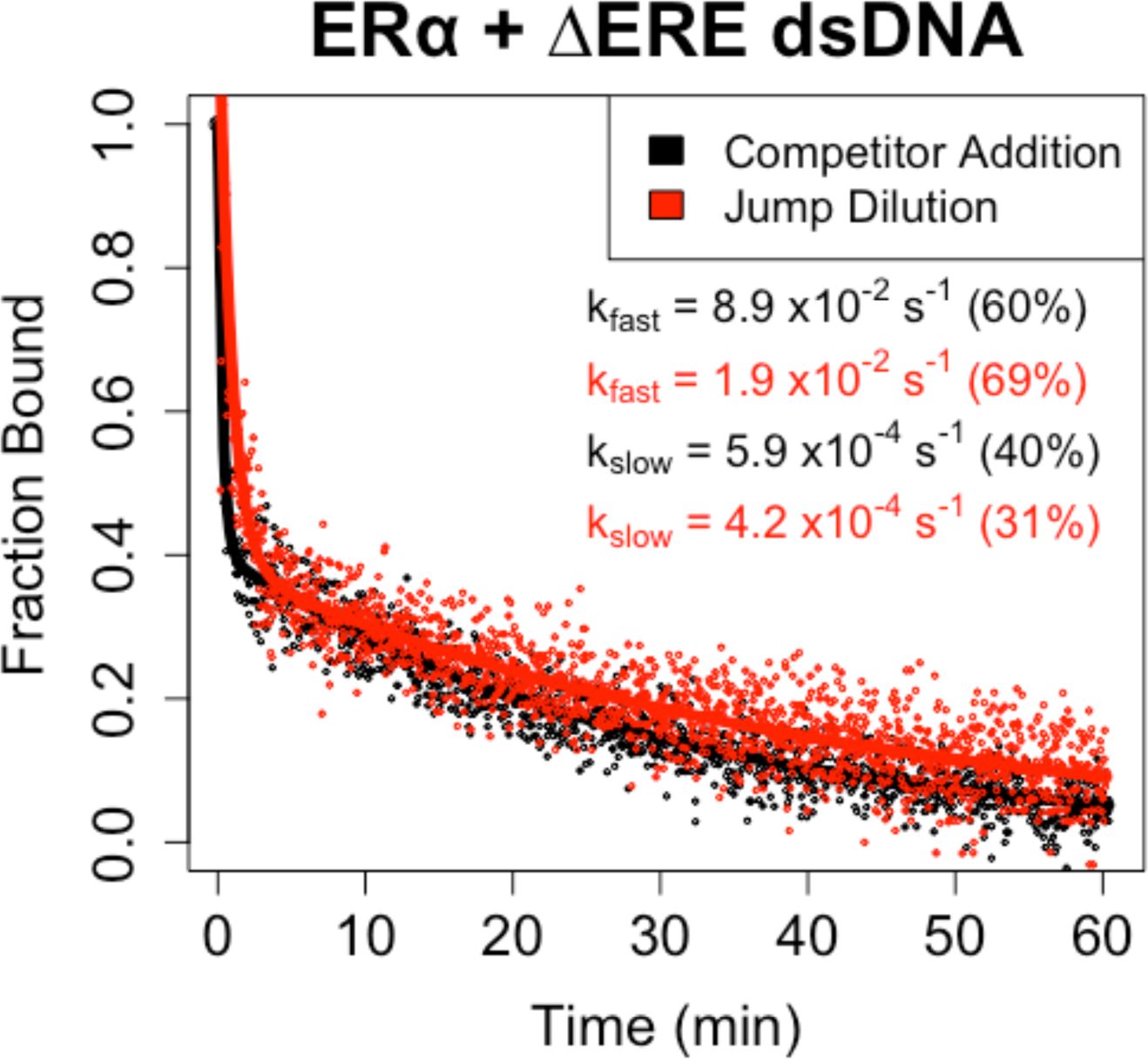
ERα_DBD-Ext_ Multiphasic Ligand Dissociation is Independent of Competitor. *Dissociation curves from FPCD versus FPJD experiments.* FPJD experiments were performed for the ERα_DBD-Ext_-ΔERE interaction using 50 nM ligand and 50 nM protein (pre-dilution), with protein-ligand reactions being incubated to equilibrium before dilution. The protein-polynucleotide reaction was then diluted ∼80-fold in buffer (at 4°C) and polarization was monitored over time post-dilution (at 4°C) to quantify protein-ligand dissociation kinetics. FPCD experiments were performed as described for Figure 1. Anisotropy traces were normalized to the controls to give ‘Fraction Bound’ over time, then normalized data were fit with bi-exponential regression to determine rate constants. Dots are data points and solid lines are regression fits (Eq. 3) from single experiments. Rate constants are the average values from all independent experiments (2 for FPCD, 3 for FPJD), with percent contributions of fast and slow components to the bi-exponential curve in parentheses.

For ERα_DBD-Ext_, we measured ΔERE dsDNA dissociation since it had the binding properties most compatible with the limitations of FPJD methodology. Notably, ΔERE dissociation was still biphasic in the absence of competitor (Figure 2), with no apparent reduction in the contribution of the fast component to the bi-exponential regression (FPCD ≈ 60%, FPJD ≈ 69%). In essence, the biphasic nature of the dissociation curves was not attributable to direct transfer. However, the presence of competitor appeared to greatly (4-5x) increase the rate of the fast component (FPCD_fast_ ≈ 8.9×10^-2^ s^-1^ vs FPJD_fast_ ≈ 1.9×10^-2^ s^-1^), but not slow component (FPCD_slow_ ≈ 5.9×10^-4^ s^-1^ vs FPJD_slow_ ≈ 4.2×10^-4^ s^-1^), of the bi-exponential regression. This suggests that the ERα_DBD-Ext_-ΔERE complex state associated with fast dissociation is susceptible to direct transfer, while the state associated with slow dissociation is not. For Sox2_HMG_, we are unable to make a similar assessment, because the FPJD assay produces less signal-to-noise relative to the FPCD assay and the Sox2_HMG_-CBS interaction has a lower anisotropy dynamic range than the ERα_DBD-Ext_-ERE interaction. Thus, the Sox2_HMG_ dissociation curves were too noisy for reliable analysis.

Ruling out direct transfer as the origin of the biphasic kinetics, we moved on to a second hypothesis. A prior *in vitro* study demonstrated that the RNA binding affinity of our ERα_DBD-Ext_ construct is facilitated by the stretch of basic residues from the hinge region of the protein, while these residues don’t significantly affect binding affinity for ERE dsDNA (Steiner et al. 2022). Thus, we assessed whether these additional nucleic acid binding residues in the construct explain the biphasic nature of our ERα_DBD-Ext_ dsDNA dissociation kinetics by providing a lower-affinity alternative binding site. We therefore compared the dsDNA dissociation kinetics for the ERα_DBD-Ext_ construct to a construct lacking the basic hinge region residues (ERα_DBD_) by FPCD (Figure 1a). Our findings indicated that ERα dsDNA dissociation kinetics were still biphasic with ERα_DBD_, ruling out these additional basic residues as the cause for biphasic dissociation (Supplemental Figure 2).

### ERα_DBD-Ext_, but not Sox2_HMG_, exhibits multiphasic association to target DNA

The observation of biphasic ligand dissociation kinetics implies the presence of multiple complex states. To probe this further, we performed FP-based ligand association experiments for ERα_DBD-Ext_ binding to ERE, ΔERE, and XBP1, and for Sox2_HMG_ binding to CBS and rG4 (Figure 3a-e). Sox2_HMG_ binding to CBS dsDNA was strictly monophasic (Supplemental Figure 3d). Sox2_HMG_ rG4 association appeared classically biphasic (i.e., fitting a bi-exponential) (Supplemental Figure 3e), where the second Sox2_HMG_ rG4 association phases in the bi-exponential regressions were ∼15x slower than the first phases. We note, however, that percent slow association phase contributions trended downward from ∼35% at 1 µM protein to <10% at 8 nM protein (Supplemental Figure 3e), resulting in monophasic association at lower protein concentrations. This trend seemed to correlate to the second transition in the binding curve, and it suggests a monomer-dimer equilibrium (Supplemental Figure 1b).

**Figure 3:**
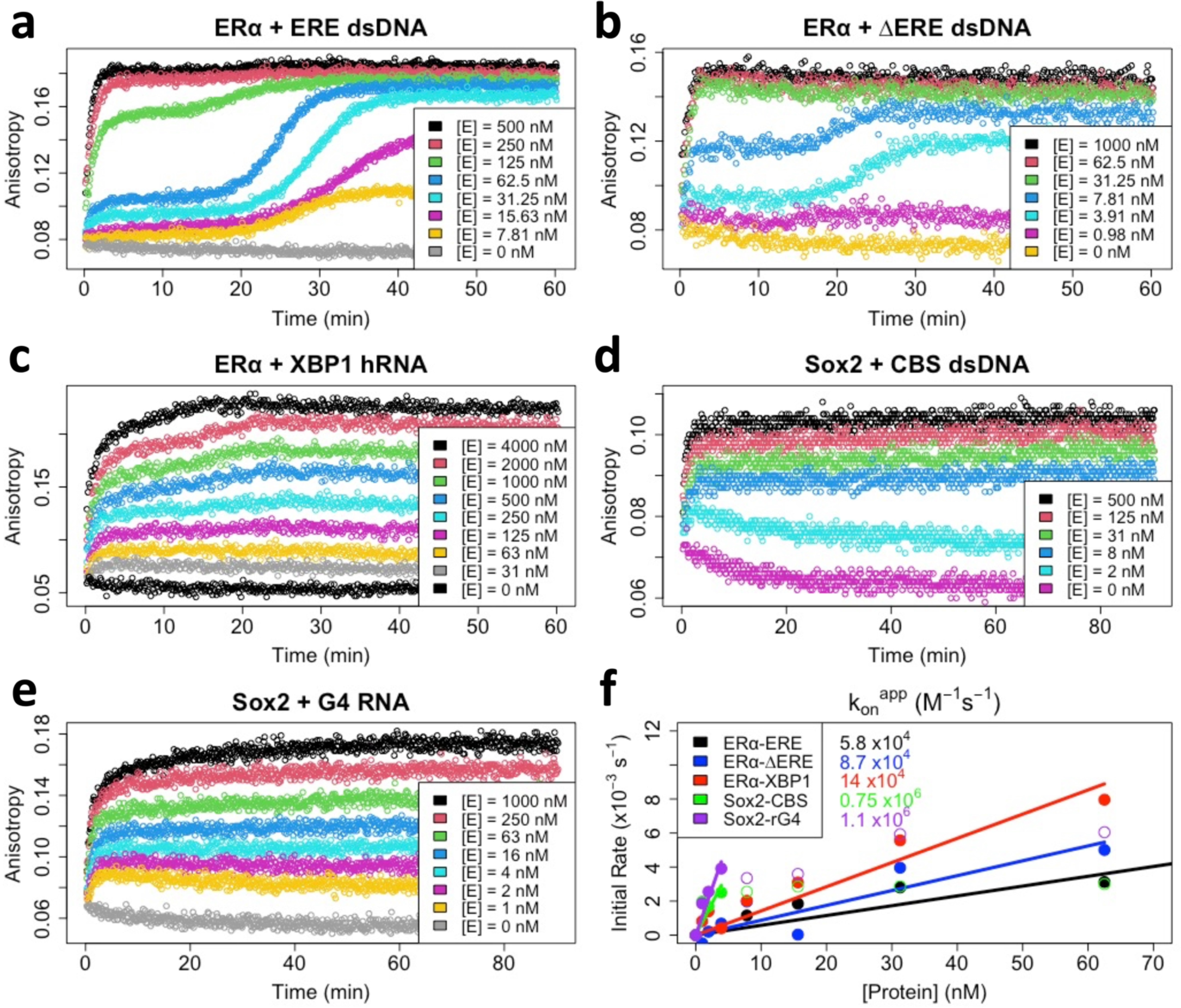
ERα_DBD-Ext_, but not Sox2_HMG_, Exhibits Multiphasic Target DNA Association. **[a-e]** *FP-based association curves.* Protein-ligand reactions were prepared after thermal equilibration (4°C), and anisotropy monitored over time immediately after protein addition to quantify association kinetics. Ligand concentrations were 5 nM, and protein concentrations ([E]) are indicated. Data are from single representative experiments (of 1-3 per protein-ligand interaction). The first 10-45 min of association data were subjected to regression with an equation for monophasic (Eq. 2.1) or biphasic (Eq. 2.2) association; the regression fits are shown in Supplemental Figure 3. **[f]** *Association rate constant analysis.* Apparent initial association rates were determined with smoothing spline regression (see Methods) and are plotted as a function of protein concentration. Each interaction has an initial linear component, followed by a plateau in apparent association rate at higher protein concentrations, which corresponds to the incomplete association curves seen at higher protein concentrations due to methodological limitations. Apparent association rates from these linear stages were used for 0-intercept linear regression to calculate apparent association rate constants (k_on_^app^). Filled circles are data points used for linear regression, open circles are data points excluded from linear regression, and solid lines are linear regression fits. Rate constants are reported with error in Table 1.

In contrast, ERα_DBD-Ext_ exhibited multiphasic association for both DNA and RNA (Figure 3a-c). For dsDNA association, we observed a highly unusual triphasic association that was protein concentration dependent. An approximately monophasic association phase was complete in ∼2 minutes (Supplemental Figure 3a-b), followed by a 10-20 minute “lag” phase, and ending in a second association phase that completes on a similar time scale as the first association phase (Figure 3a-b). Perplexingly, the “lag” phase was only evident at protein concentrations ≤10xK_d_ of the DNA, while at higher concentrations ERα_DBD-Ext_ DNA association appeared monophasic (Figure 3a-b). For ERα_DBD-Ext_ hRNA association, we observed what appeared to be biphasic association, but regression with a bi-exponential equation revealed an inadequate fit (Supplemental Figure 3c). On closer inspection, the ERα_DBD-Ext_ hRNA association curves are more like the ‘transition’ protein concentrations in the ERα_DBD-Ext_ dsDNA association curves (e.g., Figure 3a, [E] = 125 nM), suggesting similar association mechanisms.

We used the first phases of the association curves to determine apparent initial association rate constants (Figure 3f and Table 1). For ERα_DBD-Ext_, we find that initial nucleic acid association is ∼7,000-17,000x slower than diffusion-limited binding (Table 1) (Fersht 1985). For Sox2_HMG_, we find that CBS dsDNA and G4 RNA initial associations are likewise slower than diffusion-limited binding (∼1,000-1,300x) (Table 1). These rates are notably slower than some TFs, suggesting a potential conformational barrier during initial ERα_DBD-Ext_ nucleic acid binding (Halford and Marko 2004).

To gain insights into the underlying ERα_DBD-Ext_ nucleic acid binding mechanism(s), we compared our measured rate constants with the equilibrium binding data. We noted that ERα_DBD-Ext_ associates at a similar or modestly greater rate with hRNA versus dsDNA (Figure 3f), despite having lower affinity and similar dissociation rates (Figure 1b and Supplemental Figure 1a). Dividing the k_off_^app^ (apparent dissociation rate constants) by the k_on_^app^ for the respective ligands, which should yield the K_d_^app^, suggests that the ERα_DBD-Ext_ dsDNA K_d_^app^ is mostly influenced by the slow bi-exponential dissociation phase (k_off_^app^_slow_ / k_on_^app^; <u>ERE = 13 nM</u>, <u>ΔERE = 6.8 nM</u>, XBP1 = 4.7 nM) while the RNA K_d_^app^ is mostly influenced by the fast bi-exponential dissociation phase (k_off_^app^_fast_ / k_on_^app^; ERE = 0.89 µM, ΔERE = 1.0 µM, <u>XBP1 = 400 nM</u>). We made similar comparisons for the Sox2_HMG_ CBS and rG4 interactions (k_off_^app^_slow_ / k_on_^app^ and k_off_^app^_fast_ / k_on_^app^; CBS = 1.7 and 67 nM, rG4 = 0.87 and 49 nM). These values suggest that the Sox2_HMG_ CBS and rG4 fast versus slow dissociation phases (Figure 1c) could correspond to the complex states in the low versus high affinity binding curve transitions (Supplemental Figure 1b), respectively. We note for the Sox2_HMG_ G4 RNA interaction that association was biphasic (Supplemental Figure 3e), while the use of k_on_^app^ in these calculations corresponds to the initial association phase only.

### ERα_DBD-Ext_ multiphasic dissociation is not due to a “locked” binding conformation

We then sought a molecular model to explain the multiphasic dissociation kinetics observed for ERα_DBD-Ext_. An *in vitro* study of the full-length glucocorticoid receptor (GR), a nuclear hormone receptor with strong similarities to ERα, reported multiphasic dsDNA dissociation kinetics remarkably similar to the ERα_DBD-Ext_ dsDNA dissociation kinetics observed here (De Angelis et al. 2015). Those authors proposed a “locked” binding conformation model to explain GR multiphasic ligand dissociation kinetics (Supplemental Figure 4a). This model suggests that after initial GR-dsDNA association, the complex can slowly isomerize to an alternative state but must slowly isomerize back to the initial complex state before ligand dissociation can occur. A prediction of this model is that if a brief protein-ligand incubation period is allowed before complex dissociation is induced (e.g., by competitor addition), then the complex should not have time to isomerize to the more stable alternative state, and the slow phase of the dissociation kinetics should be ablated.

To test if ERα_DBD-Ext_ ligand dissociation kinetics could be explained by the “locked” binding conformation model, we conducted FPCD experiments with variable protein-ligand incubation times. Our findings indicated that 2-min versus 60-min protein-ligand incubations produced similarly biphasic ERα_DBD-Ext_ dsDNA dissociation, with slow phase contributions of 30-40% based on biexponential regression (Supplemental Figure 4a-b). In contrast, based on our Figure 1 data, the GR model predicts a ∼5% slow phase contribution after a 2-min incubation. The behavior was somewhat different when RNA was the ligand – the shorter incubation time did affect ERα_DBD-Ext_ hRNA dissociation, but by partially reducing the fast phase of the bi-exponential regression instead of the anticipated slow phase reduction (Supplemental Figure 4c). It’s notable that the ERα_DBD-Ext_ hRNA association is incomplete after a 2-min incubation at the protein concentrations used (Supplemental Figure 3c), suggesting that the fast dissociation phase of the bi-exponential regression emerges during the second ERα_DBD-Ext_ hRNA association phase (Figure 3a-c). Overall, despite the similarities in dissociation kinetics, our data indicate that the previously proposed model for GR multiphasic ligand dissociation does not apply to ERα_DBD-Ext_ DNA or RNA biphasic dissociation.

While these findings were sufficient to refute one model, we further investigated how ERα_DBD-Ext_ dsDNA complex stability varied during its multiphasic association to provide insights into alternate models. The above dsDNA experiments used ERα_DBD-Ext_ concentrations several fold above the ligand K_d_, where nucleic acid association occurs in a single apparent step (Figure 3a). To determine if protein-ligand incubation time affects dissociation kinetics at lower protein concentrations, when association is multiphasic, we performed FPCD experiments under these conditions using ERE dsDNA (Figure 4). We selected protein-ligand incubation times (Figure 4a for reference) just after initial association at the beginning of the “lag” phase (2.5 min), at the end of the “lag” phase before secondary association (15 min), towards the end of secondary association (30 min), and at binding equilibrium (60 min). These findings indicate that ERα_DBD-Ext_ ERE dsDNA dissociation is slow and monophasic during the “lag” phase after initial association, but complex dissociation acquires a faster component and becomes biphasic during the second association phase (Figure 4b-e). Curiously, this suggests that the more stable complex state emerges first, followed by the less stable complex state, which contrasts with the positive cooperativity observed by the ERα_DBD-Ext_ ΕRΕ binding curve (Supplemental Figure 1a). Notably, this is the same trend in dissociation behavior over multiphasic association that was observed for the ERα_DBD-Ext_ XBP1 hRNA interaction above, consistent with the hypothesis that the two ligands could share an underlying mechanism.

**Figure 4:**
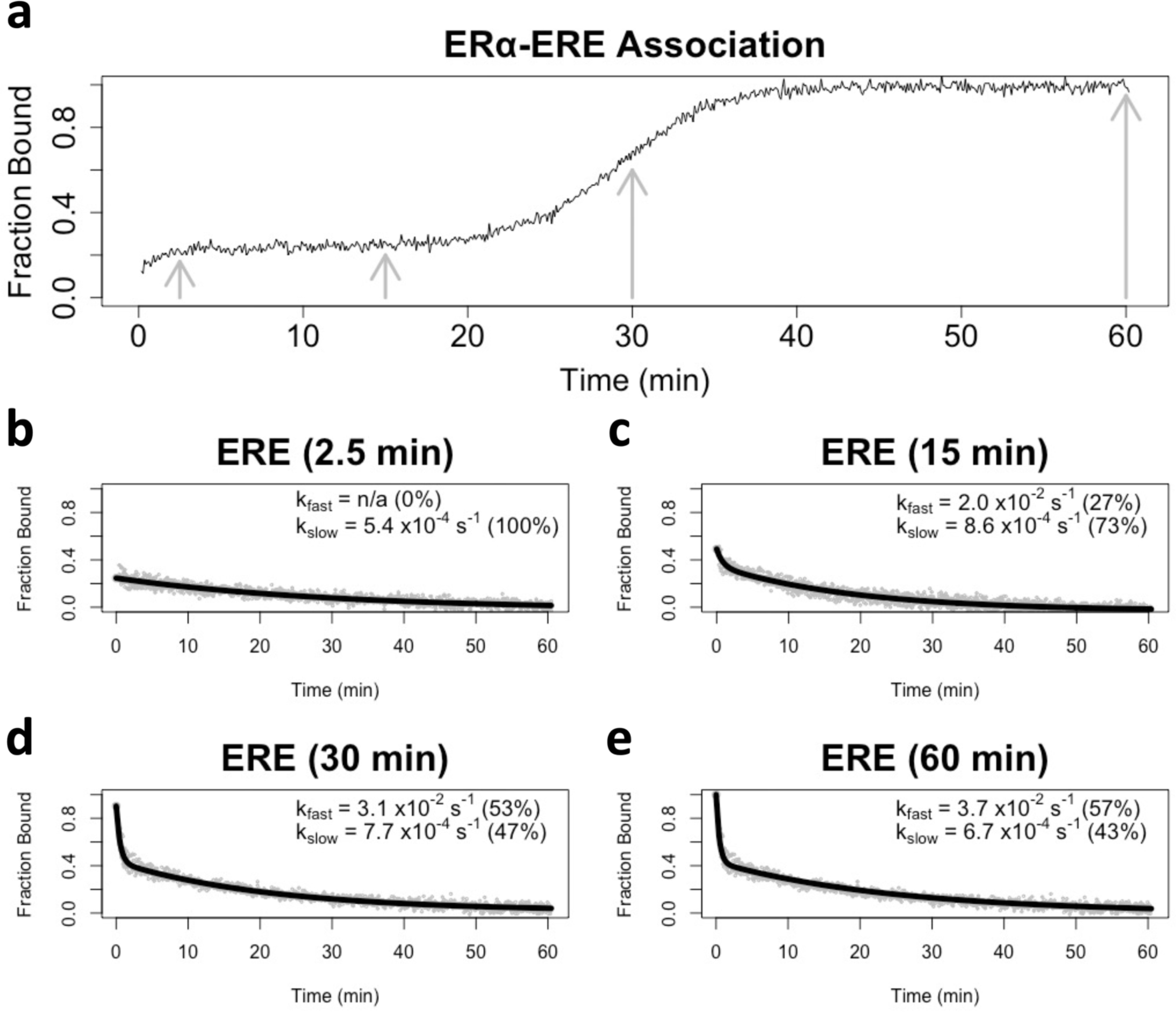
The More Stable Complex State Forms First During Multiphasic ERα_DBD-Ext_ Ligand Association. **[a]** *Normalized ERα_DBD-Ext_ association curve.* Normalized association curve of 30 nM ERα_DBD-Ext_ and 2 nM ERE dsDNA, taken from Figure 3a. Grey arrows correspond to incubation times prior to competitor addition in panel b-e experiments. **[b-e]** *Dissociation curves from FPCD experiments where competitor was added after variable protein-ligand incubation times.* FPCD experiments (see Figure 1a) were performed for the ERα_DBD-Ext_-ERE interaction using 2 nM ligand, 10 µM competitor, and 30 nM protein; protein-ligand reactions were incubated for 2.5 min (b), 15 min (c), 30 min (d), or 60 min (e) before competitor addition. Anisotropy traces were normalized to the internal controls to give ‘Fraction Bound’ over time, then normalized data were fit with bi-exponential regression (Eq. 3) to determine rate constants. Grey dots are data points and solid black lines are regression fits from single experiments. Percent contributions of fast versus slow components to the bi-exponential curve are in parentheses.

### ERα_DBD-Ext_ multiphasic DNA dissociation is conserved across methodology and temperature

To ensure that our findings were not due to an unexpected feature of our FP experimental design, we used surface plasmon resonance (SPR) to independently measure ERα_DBD-Ext_ dsDNA association and dissociation kinetics (Supplemental Figure 5). This also provided the opportunity to obtain data at a second temperature (25°C vs 4°C). Given our assay requirements and the limitations of SPR, we could only measure the association kinetics up to 5 minutes, which is before the secondary association phase that emerged in FP assays. Using SPR association curves to calculate ERα_DBD-Ext_ apparent association rate constants for ERE and ΔERE dsDNA (Figure 5b), we infer that they are 2-3x higher than the respective values determined by FP (Table 1), which puts them in good agreement given the higher temperature for SPR experiments.

**Figure 5:**
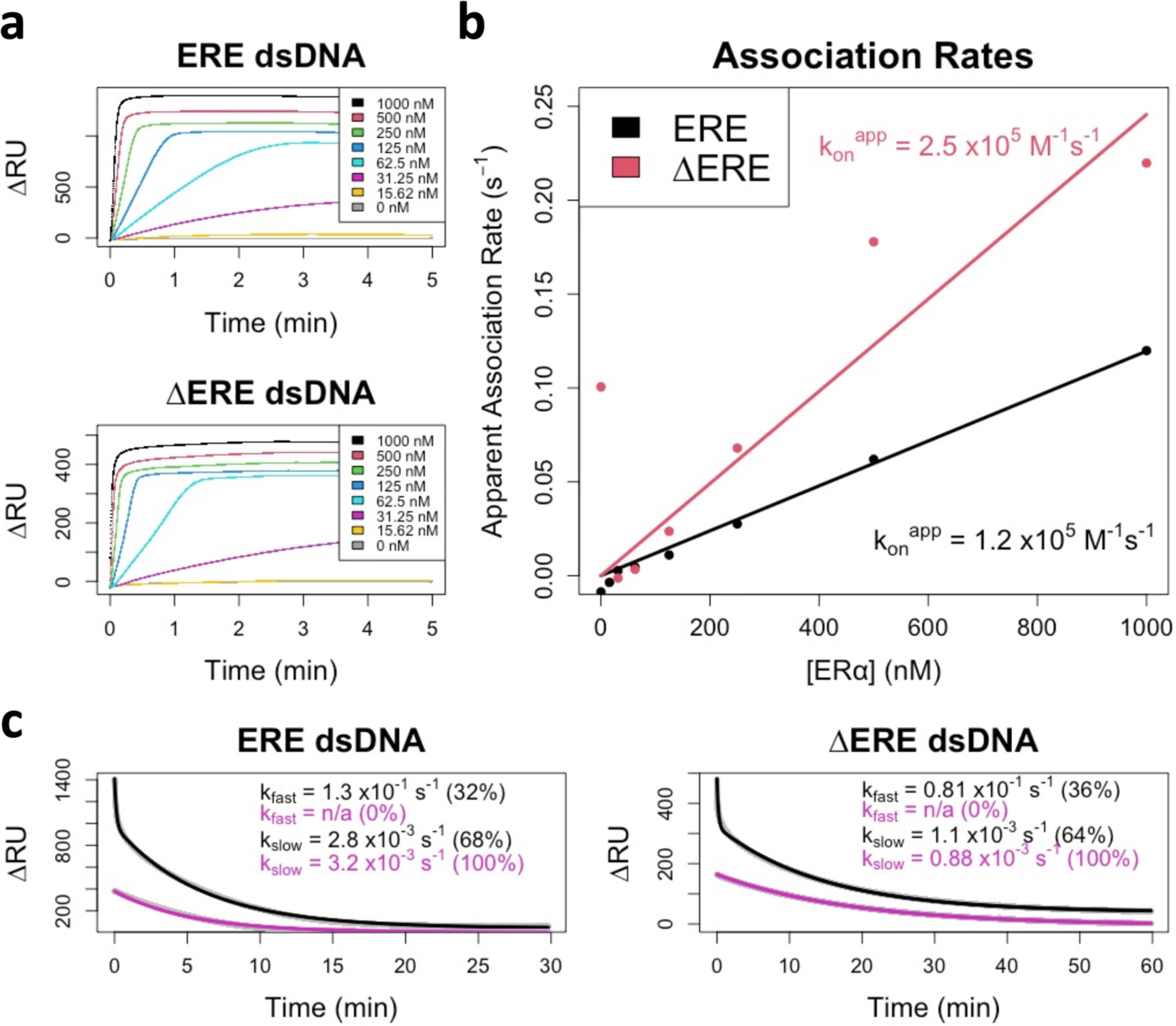
Surface Plasmon Resonance Confirms Multiphasic ERα_DBD-Ext_-dsDNA Binding Kinetics. **[a]** *ERα_DBD-Ext_ ligand association curves.* SPR association phase curves, taken from Supplemental Figure 5. Legends indicate the concentration of ERα_DBD-Ext_ used during protein injection. Lines are data, not regression fits. **[b]** *Association rate analysis.* Association curves from panel a had their initial slopes normalized to their signal dynamic range to calculate their apparent association rates (see Methods). Plots of apparent initial association rates versus protein concentrations were fit with 0-intercept linear regression to calculate apparent initial association rate constants (k_on_^app^). Dots are data and solid lines are linear regression fits. **[c]** *Dissociation rate analysis.* SPR dissociation phase curves, taken from Supplemental Figure 3. Two key protein concentrations (1 µM and 30 nM) are shown, corresponding to the color scheme in panel a. Data were fit with bi-exponential regression (Eq. 3) to determine rate constants. Grey dots are data and solid black/purple lines are regression fits; data points are mostly obscured by regression lines. Percent contributions of fast versus slow components to the bi-exponential curve are in parentheses. All SPR data in this figure are from a single experiment per ligand.

Based on the FP data, we predicted that SPR could be used to test the dissociation behavior at different protein concentrations. As noted above, SPR couldn’t accommodate the time range to fully repeat the FP experiments that revealed multiphasic association (Fig 3) or dsDNA dissociation kinetics over the time course of its multiphasic association (Fig 4). However, we estimated that complexes formed at lower ERα_DBD-Ext_ concentrations should begin their dissociation curves in the “lag” phase of multiphasic association, while complexes formed at higher protein concentrations should begin their dissociation curves at equilibrium. As predicted, for both ERE and ΔERE dsDNA the ERα_DBD-Ext_ dissociation curves were biphasic at high ERα_DBD-Ext_ concentrations but monophasic slow at low ERα_DBD-Ext_ concentrations (Figure 5c), suggesting that our SPR and FP findings are reporting the same phenomenon. The SPR-derived rate constants for the fast and slow phases of the bi-exponential regressions (Fig. 5c) were 1-3x greater than their respective FP values (Table 1), which is in good agreement given the temperature differential. We note that the percent contribution of the fast phase to the bi-exponential regression was lower for SPR (32-36%) than for FP (60-71%). However, unlike the FP experiments (Supplemental Figure 1a), the SPR signal had not yet appeared to plateau at the highest protein concentrations, and the percent contribution of the fast phase to the bi-exponential regression still appeared to be increasing with protein concentration (Supplemental Figure 5) suggesting that these values may be more similar at saturating protein concentrations. Collectively, these SPR-based findings independently confirm the kinetic observations for our FP experiments.

## DISCUSSION

ERα and Sox2 have been previously demonstrated to bind RNA with structural specificity *in vitro*, and to associate with RNA *in vivo* (Holmes et al. 2020; Hamilton et al. 2022; Ng et al. 2012; Xu et al. 2021). In addition, their RNA and DNA interactions are reportedly competitive, but initial studies also suggest that the RNA and DNA binding surfaces do not perfectly overlap on the TFs (Steiner et al. 2022; Holmes et al. 2020; Hou et al. 2020). Our work expands on these *in vitro* findings by elucidating the timescales for ERα and Sox2 nucleic acid association and dissociation, and by interrogating their respective RNA versus DNA binding mechanisms.

### A model for Sox2_HMG_ DNA and RNA binding

Our kinetic and thermodynamic data allow us to propose a minimum kinetic model for Sox2_HMG_ binding to nucleic acids. Binding to target (CBS) dsDNA and G4 RNA exhibited two-transition equilibrium binding (Supplemental Figure 1b), biphasic dissociation (Figure 1c), and monophasic dsDNA association and biphasic G4 RNA association (Figure 3d-e, Supplemental Figure 3d-e). The second, lower affinity Sox2_HMG_ CBS and rG4 binding transitions have similar affinities to non-target DNA and RNA. Prior data indicate that Sox2 _HMG_ can bind DNA and RNA at protein-ligand stoichiometries higher than 1:1 (Holmes et al. 2020; Hamilton et al. 2022; Moosa et al. 2018). We propose that the simplest model to sufficiently explain these findings is a sequential protein binding model (Figure 6a). We also considered a ligand isomerization (but not protein isomerization) model as shown in Figure 6b that would be consistent with the data if the ligand states were in comparable proportions at equilibrium and had drastically different affinities for the protein. However, single dominant bands were observed via native-PAGE during nucleic acid preparation (see methods), no two-transition binding curves were produced for other dsDNA ligands (Supplemental Figure 1b), and G4 RNAs are normally quite stable *in vitro* under our 135 mM KCl conditions (Lane et al. 2008; Crenshaw et al. 2015). Consequently, we don’t favor the Fig. 6b ligand isomerization model.

**Figure 6:**
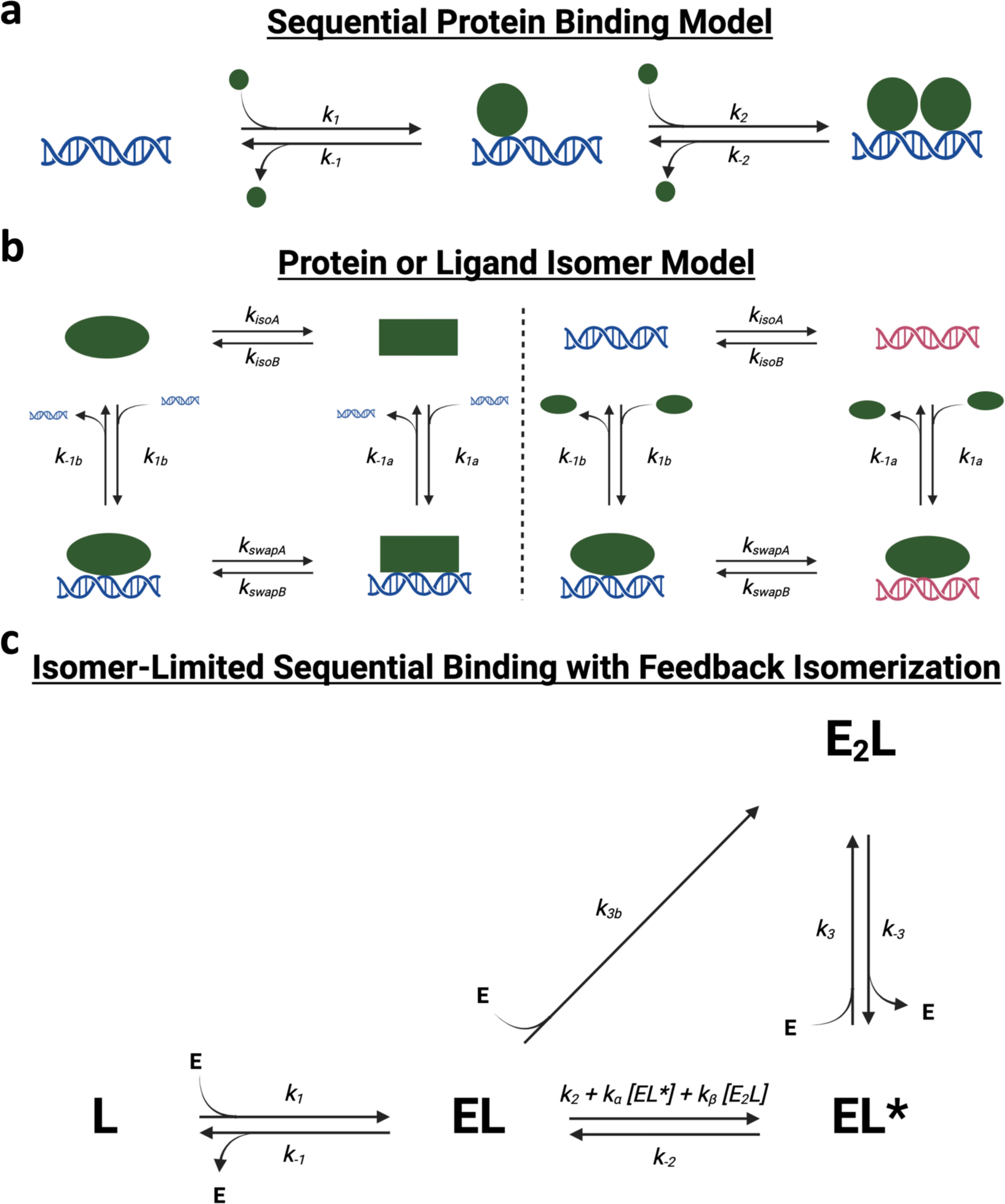
Various Reaction Schemes that Predict Biphasic Protein-Ligand Dissociation. **[a]** *Sequential Protein Binding Model.* After initial protein-ligand association, additional protein can associate with the complex to form higher stoichiometry complexes. If the complex states with differing stoichiometries also have differing stabilities, multiphasic dissociation kinetics can be produced. If protein associates at differing rates with ligand versus existing complex, multiphasic association kinetics can be produced. Based on this model, rate constants for Sox2_HMG_ CBS dsDNA binding are inferred to be k_1,2_ ≈ 8×10^5^ M^-1^s^-1^, k_-1_ ≈ 1×10^-3^ s^-1^, and k_-2_ ≈ 5×10^-2^ s^-1^. The rate constants for Sox2_HMG_ G4 RNA binding are inferred to be k_1_ ≈ 1×10^6^ M^-1^s^-1^, k_2_ ≈ 7×10^4^ M^-1^s^-1^, k_-1_ ≈ 1×10^-3^ s^-1^, and k_-2_ ≈ 5×10^-2^ s^-1^. **[b]** *Protein or Ligand Isomer Model.* The protein (left) or ligand (right) may isomerize to an alternative state, which produces different protein-ligand association and/or dissociation rates. **[c]** *Isomer-Limited Sequential Binding with Feedback Isomerization.* After initial protein-ligand association to form a stable complex (EL), protein can inefficiently associate with the initial complex to form higher-order stoichiometry complex (E_2_L), or the initial complex can isomerize to an alternative complex state (EL*) that can more readily accommodate additional protein monomers. Complex isomerization is intrinsically slow but may be accelerated by “feedback” from isomerized complex or higher-order stoichiometry complex. Reaction scheme could produce monophasic, biphasic, or “lagged” triphasic association, and monophasic, biphasic, or triphasic dissociation, depending on specific values of rate constants.

Under a sequential protein binding model (Figure 6a, see legend for inferred rate constants), Sox2_HMG_ initially binds target dsDNA or G4 RNA with high affinity, then an additional Sox2_HMG_ monomer (or more) binds with lower affinity. Prior studies attribute this to sequence-specific (or structure-specific) binding followed by nonspecific binding (Hamilton et al. 2022; Holmes et al. 2020). Upon addition of competitor, the fast dissociation phase results from the less stable monomer(s) dissociating followed by slow dissociation of the more stable monomer from the ligand. Since association appears approximately monophasic for DNA and biphasic for RNA (Supplemental Figure 3d-e), we infer that the DNA association rate constants for successive monomers are comparable, but RNA association rate constants decrease with sequential monomer association. Finally, we note that the single-transition Sox2_HMG_ binding curves for non-target DNA and RNA support this model – without the tight binding, Sox2_HMG_ would only bind weakly with increasing K_d_ for each subsequent monomer. Such behavior could produce the low Hill coefficients seen for nontarget DNA and RNA (Supplemental Figure 1b), and it would explain the similar affinities for the nontarget nucleic acids’ K_d_^app^ and the target DNA and RNA second transition K_d_^app^.

### A model for ERα_DBD-Ext_ DNA and RNA binding

Synthesis of our thermodynamic and kinetic data allows us to consider several models for DNA and RNA binding by ERα_DBD-Ext_. ERα_DBD-Ext_ equilibrium binding exhibited a single-transition with apparent positive cooperativity for ERE dsDNA but not for ΔERE dsDNA or XBP1 hRNA (Supplemental Figure 1a). Existing crystal structures of ERα_DBD_ binding to ERE dsDNA reveal that one ERα_DBD_ monomer binds each of the two repeats in the palindromic recognition sequence and the two monomers then stabilize one another on DNA through protein-protein interactions (Schwabe et al. 1993), which explains the apparent positive cooperativity and suggests 2:1 protein-DNA stoichiometry. By extension, it seems reasonable that the loss of one recognition sequence repeat in the ΔERE dsDNA would reduce stoichiometry and cooperativity, consistent with our data (Supplemental Figure 1a). The ERα_DBD-Ext_ dsDNA and RNA interactions also exhibited biphasic dissociation with similar rate constants (Figure 1b), and for dsDNA this persisted even in the absence of competitor (Figure 2) or hinge residues (Supplemental Figure 2). ERα_DBD-Ext_ dsDNA association was apparently triphasic at lower protein concentrations, with a 10-20 min “lag” between two typical association phases, while RNA association seemed biphasic but did not adequately fit a standard bi-exponential (Figure 3a-b). Most notably, while the ERα_DBD-Ext_ dsDNA K_d_^app^ was primarily predicted by the k_off_^app^_slow_, the RNA K_d_^app^ was more influenced by the k_off_^app^_fast_.

This raises the question of what model best explains the ERα binding data. First, a standard sequential protein binding model for biphasic dissociation (Figure 6a), which was the favored model for Sox2_HMG_ binding, is inconsistent with our data. Specifically, we note that this model can only explain biphasic dissociation from a state of saturated ligand binding if subsequent ERα_DBD-Ext_ monomers bind DNA/RNA much less stably than the first monomer, but ERα_DBD-Ext_ binding exhibits neither two-transition binding nor a low Hill coefficient (i.e., n<1) (Supplemental Figure 1a). Furthermore, such a model would not be able to produce the “lagged” triphasic association curves that we observed, even if a slow isomerization step was included between sequential monomer binding events is Figure 6a (this would produce classic biphasic association). Next, the “locked” binding conformation model (Supplemental Figure 4a), which was previously proposed to explain biphasic DNA dissociation for GR (De Angelis et al. 2015), was specifically tested and discounted by our studies (Supplemental Figure 4). We also note that this model could explain monophasic association, or biphasic association if the “locked” complex conformation has significantly altered anisotropy, but it cannot explain our “lagged” triphasic association data (Figure 3). Finally, protein or ligand isomerization models as shown in Figure 6b would require significantly different complex state stabilities to explain our data, but ERα_DBD-Ext_ DNA/RNA binding exhibits neither two-transition binding nor a low Hill coefficient (Supplemental Figure 1a). In addition, the Fig. 6b models could be reconciled with monophasic or biphasic association, but they cannot explain the “lagged” triphasic association we observed (Figure 3). We considered if heterogeneity in ligand or protein could explain the data, but the RNA/DNA had a single dominant band via native-PAGE (see Methods) and no protein heterogeneity was observed during size-exclusion chromatography or SDS-PAGE (see Methods).

None of the above models (Figure 6a-b, Supplemental Figure 4a) could explain the “lagged” triphasic association we observed (Figure 3). In contrast, one kinetic phenomenon that can produce an apparent association lag followed by seemingly rapid/spontaneous association involves sequential reactions where a downstream product has a “feedback” effect to catalyze an earlier step in the reaction. In the case of ERα_DBD-Ext_ dsDNA binding, such a minimum kinetic model might resemble that shown in Figure 6c. In this model, ERα_DBD-Ext_ initially binds dsDNA with high affinity to produce a stable complex. We note that it’s possible for the initial complex to have 2:1 instead of the 1:1 protein-ligand stoichiometry shown without drastically altering apparent kinetics. Additional ERα_DBD-Ext_ monomers would be capable of inefficiently associating with the initial stable complex, but the initial complex could also slowly isomerize to an alternate complex state that better accommodates additional ERα_DBD-Ext_ monomer binding. Critically, this complex isomerization would have to be susceptible to acceleration by already isomerized (and/or higher order stoichiometry) complex. Our simulations with the Figure 6c model suggest that it can recapitulate the kinetic trends observed for ERα_DBD-Ext_ dsDNA binding (Supplemental Figure 6, see legend). Notably, accelerating the reverse isomerization (i.e., increasing k_-2_) in simulations made the data look more like the ERα_DBD-Ext_ RNA binding kinetics (Supplemental Figure 7, see legend). In essence, our observations that ERα_DBD-Ext_ DNA and RNA binding had similar dissociation rates and initial association rates despite differing K_d_^app^, Hill coefficients, and association curve shapes could be recapitulated by this model by changing a single rate constant value.

Overall, we present evidence that ERα_DBD-Ext_ DNA and RNA binding cannot be adequately explained by traditional reaction schemes. Instead, we provide a ‘framework’ model that can generally recapitulate ERα_DBD-Ext_ DNA and RNA kinetic trends. We note that our few simulations with this model are certainly not a perfect fit to the experimental data herein. Indeed, the “flexibility” of the model made it difficult to exhaustively fit our experimental data via iterative numerical integration and regression. Thus, it’s likely that our Figure 6c model does not completely capture the mechanism(s) of ERα_DBD-Ext_ DNA and RNA binding. Rather, the model represents a starting point for insights into the ‘true’ mechanism. First, the model suggests that, despite the seemingly disparate ERα_DBD-Ext_ DNA versus RNA association behaviors and affinities, they could share a reaction mechanism. Second, the hallmark “lagged” triphasic association we observed is critically dependent on reaction mechanism “feedback” and complex isomerization in our simulations. While it’s not hard to imagine a TF like ERα that binds gene targets as a homodimer having more than one conformational state after target binding, the novel implication that some ERα_DBD-Ext_ nucleic acid binding states can influence the stability of other complex states warrants further investigation. Such a mechanism could be especially relevant in situations like ERα nucleic acid condensates where multiple complexes are crowded together (Nair et al. 2019), since overall condensate stability and architecture could be impacted if certain complex states influence the stabilities of other complex states and their ability to accommodate additional protein monomers.

### Concluding remarks

In a cell, hundreds to thousands of transcription factors search for their unique target DNA sequences to perform critical regulation of gene expression. During this target search, the TFs not only coordinate with many protein binding partners, but they are also inundated with numerous other nucleic acids like the nontarget DNA in surrounding chromatin and free and nascent RNA. Consequently, TFs likely experience a variety of transitory nucleic acid binding events on the way to their target DNA sites. This is likely to be even more pronounced in the dense environment of biological condensates. Discerning the relevance of a TF’s numerous nucleic acid interactions to its biological function requires careful consideration of the prevalence, lifetimes, reaction mechanism(s), and inter-ligand influences of these varied binding events. Our biophysical studies herein estimate the timescales of association and dissociation events for the RNA and DNA interactions of two model TFs, and they also provide insight into the reaction mechanism(s) for these TF nucleic acid interactions. Thus, our findings represent a valuable ‘touchstone’ for considerations of how ERα, Sox2, and other related TFs have their target site searches influenced by competing nucleic acids like RNA.

## MATERIALS & METHODS

### Protein Expression and Purification

For recombinant Sox2_HMG_, a plasmid encoding Sox2_HMG_ as an N-terminal octa-histidine and maltose binding protein (MBP) fusion with rhinovirus 3C protease cleavage site in a pET30b vector was generously provided by Desmond Hamilton (Batey lab, University of Colorado Boulder) (Hamilton et al. 2022). The plasmid was transformed into BL21(DE3) *E. coli* bacterial cells, inoculated into 20 mL media (LB broth with 100 µg/mL kanamycin), then the starter culture incubated overnight at 37°C/200rpm until A_600_ ≈ 5.0. The culture was then induced with 0.5 mM Isopropyl β-D-thiogalactopyranoside (IPTG) and incubated at 37°C/200rpm for an additional 4 h. Following induction, the culture was pelleted by centrifugation (4,000x g/4°C/20min) and resuspended in 50 mL Amylose A buffer (20 mM TRIS pH 7.5 at 25°C, 200 mM NaCl, 1 mM EDTA) with 1 Pierce Protease Inhibitor Tablet (Thermo Scientific #A32965) and 50 mg lysozyme (Sigma Aldrich #L6876), then lysed with an Emulsiflex C3 homogenizer (Avestin) at 15,000-18,000 psi. Lysate was clarified by centrifugation (27,000x g/4°C/30min) and the supernatant collected. A 4°C AKTA Pure FPLC system (Cytiva) was prepared with a 10 mL amylose column (NEB #E8021S) and 2 mL/min flow rate, equilibrated with Amylose A buffer, supernatant applied, washed with Amylose A buffer, and eluted with Amylose B buffer (20 mM TRIS pH 7.5 at 25°C, 200 mM NaCl, 1 mM EDTA, 10 mM maltose). To the eluent was added 1.0 mg of Prescission Protease, and I was then loaded into 10 kDa-cutoff SnakeSkin Dialysis Tubing (Thermo Scientific #68100) and dialyzed overnight at 4°C in P-cell A buffer (50 mM TRIS pH 7.5 at 25°C, 1 mM EDTA) with 50 mM NaCl. The FPLC system was prepared with a 10 mL P11-phosphocellulose column (Whatman) and 2 mL/min flow rate, equilibrated with P-cell A buffer, dialyzed eluent applied, washed with P-cell A buffer, and eluted with a 100 mL 0-100% P-cell B buffer (50 mM TRIS pH 7.5 at 25°C, 1 mM EDTA, 1 M NaCl) gradient. Protein-containing (via A_280_) fractions were reconciled and concentrated with a 10 kDa-cutoff centrifugal filter unit. The FPLC system was prepared with a Superose 6 size-exclusion column and 0.25 mL/min flow rate, then equilibrated with Sizing Buffer (10 mM TRIS pH 7.5 at 25°C, 250 mM NaCl, 1mM EDTA), concentrated eluent applied, and followed with Sizing Buffer. Protein-containing (via A_280_) eluent fractions were reconciled for final product, then flash frozen in liquid nitrogen and stored at -80°C. SDS-PAGE indicated ≥95% purity, and protomer concentrations were determined by spectroscopy with ε_280_ = 13,980 M^-1^cm^-1^. One liter of culture typically yielded ∼4 mg of final protein.

For recombinant ERα_DBD_ and ERα_DBD-Ext_ (residues 180-262 for ERα_DBD_,180-280 for ERα_DBD-Ext_), proteins were expressed with a thrombin-cleavable N-terminal hexahistidine tag using a pET28a (EMD Biosciences) vector. Protein expression and purification methods were described previously (Steiner et al. 2022). Starting with a single transformed colony of BL21(DE3)pLysS *E. coli*, expression cultures were grown at 37°C (with 50 μg/mL kanamycin and 50 μg/mL chloramphenicol) using 2x YT rich media to an OD_600_ of 0.8-1.0 and cold shocked on ice for 20 min. IPTG was added to a final concentration of 0.4 mM, along with 50 μM ZnCl_2_, to induce protein expression and cultures were grown for 3 h at 37°C in a shaker. Cells were harvested by centrifugation (5,000 x g) and pellets stored at −20°C. Cell pellets were thawed and resuspended in 50 ml lysis buffer (20 mM Tris pH 7.5 at 25°C, 1 M NaCl, 10 mM imidazole pH 7.5, 5% glycerol) per 1 L of cells with one EDTA-free protease inhibitor cocktail tablet (Roche). Cells were lysed using a Misonix Sonicator 3000 (110 W for 2 min total ON-time, pulse: 15 s ON/45 s OFF, ½ inch tip) and the lysate cleared by centrifugation (15,000x g, 30 min). Cleared lysate was loaded onto lysis buffer-equilibrated Ni-NTA resin (GoldBio, 5 mL resin per 50 mL lysate) and rocked gently for 1 h at 4°C. The bead slurry was loaded onto a gravity flow column and washed twice with increasing concentrations of imidazole in lysis buffer (wash 1: 20 mM imidazole, wash 2: 30 mM imidazole), then eluted with 300 mM imidazole in lysis buffer. Bovine α-Thrombin (Haematologic Technologies Incorporated) was added (10 U/mg protein) to the eluate to remove the hexahistidine tag. The eluate solution was transferred to 6-8 kDa MWCO dialysis tubing (Spectra/Por – Spectrum Labs) and dialyzed overnight at 4°C in 4 L of column buffer (20 mM Tris pH 7.5 at 25°C, 100 mM NaCl, 5% glycerol, 1 mM DTT). Dialyzed eluate was filtered to 0.2 μm and concentrated using 5 kDa MWCO spin concentrators (Vivaspin Turbo). The sample was again filtered to 0.2 µm and loaded onto a HiLoad 16/600 Superdex 75 column (GE Healthcare) and eluted as a monomer. Pooled fractions containing recombinant ERα were assessed for purity, aliquoted, flash-frozen, and stored at −70°C. One liter of culture typically yielded 2 mg of purified protein as measured by absorption (ε_280_ = 14,440 M^−1^cm^−1^). All experiments used ERα_DBD-Ext_ protein, unless otherwise indicated.

### Preparation of Oligonucleotides

All oligonucleotides except the XBP1 hRNA were ordered from IDT (Coralville, IA), and their sequences in IDT syntax are provided (Supplemental Table 1). For Sox2_HMG_ dsDNA ligands, complementary oligonucleotides ordered from IDT were mixed at 100 µM each in annealing buffer (50 mM TRIS pH 7.5 at 25°C, 200 mM NaCl) and subjected to a thermocycler program (95°C for 10 min, 95→4°C at 0.5 °C/min, hold at 4°C) for annealing. For ERα dsDNA, the complementary strands were combined at 1 μM labeled and 5 uM unlabeled (ligand prep) or 100 µM each unlabeled (competitor prep) in annealing buffer (20 mM Tris pH 7.5 at 25°C, 50 mM NaCl), then annealed by bench-top slow cooling (heated 95°C for 1 min, then cooled to room temperature for 3 h). Complete annealing for all oligonucleotides was confirmed via native-PAGE. Concentrations of all ligands were confirmed spectroscopically using manufacturer-provided extinction coefficients.

According to prior methodology (Steiner et al. 2022), XBP1 hRNA was prepared by *in vitro* transcription (IVT) with T7 RNA polymerase, using dsDNA templates containing a T7 polymerase promoter sequence, which were created via PCR with IDT-synthesized oligonucleotides. Briefly, full PCR amplification was confirmed on 2% agarose gel, and subsequent IVTs were performed for 3 h at 37°. Successful transcription was confirmed via 10-18% denaturing PAGE. After IVT, RNAs were precipitated in 1/10^th^ volume 3M sodium acetate and 2.5 volumes ice cold ethanol overnight. The following day ethanol precipitated RNA was pelleted and dried, then resuspended and purified by denaturing urea-polyacrylamide gel electrophoresis followed by buffer exchange and concentration in a 5 kDa MWCO spin concentrator (Vivaspin Turbo). Purified RNA oligonucleotides were 3′-end labeled with pCp-AF488 (Alexa Fluor). 200 pmol of RNA and 2400 pmol fluorophore were combined in labeling buffer (1mM ATP, 10% DMSO, 50% PEG, 40 U T4 ligase and 1x T4 ligase buffer) overnight at 16 ° C. The labeled RNA was purified using RNA Clean & Concentrator kit (Zymo #4060), passed through a G-25 spin column, and stored at -20°C. Concentration was determined by A_260_ and total RNA yield was typically 10–50% with ∼70% labeling efficiency. Purity of the final sample was assessed by 10–15% denaturing PAGE and imaged by fluorescence (Ex = blue wavelength filter, Em = green wavelength filter). RNA samples were prepared for binding assays by fast refolding at 1 μM by snap-cooling (95°C for 1 min, ice for >5 min).

### FP-based K_d_^app^ and Association Rate Determination

Pre-reaction mix was prepared with 5 nM ligand in ERα binding buffer (20 mM TRIS pH 7.5 at 25°C, 100 mM NaCl, 5% glycerol, 0.01% IGEPAL) or Sox2 binding buffer (10 mM TRIS pH 7.5 at 25°C, 135 mM KCl, 15 mM NaCl, 0.1 mg/mL nonacetylated BSA, 4% Ficoll, 0.05% NP-40, 1 mM DTT), then dispensed in 36 μL volumes into the wells of a 384-well black microplate (Corning #3575). A range of protein concentrations was prepared at 10x the final reaction concentrations via serial dilution in respective binding buffer. Protein and microplates were then thermally equilibrated at 4°C for 30 min. After thermal equilibration, reactions were initiated by addition of 4 μL of the respective 10x protein concentration to the corresponding microplate well, then incubated for ≥60 min at 4°C. Fluorescence anisotropy readings were taken over the course of incubation immediately after protein addition (in <10 s intervals) with a TECAN Spark microplate reader (Ex = 481 ± 20 nm, Em = 526 ± 20 nm). Each experiment had 1 reaction (well) per protein concentration, and 1-3 independent experiments were performed per protein-polynucleotide interaction.

For equilibrium dissociation constant calculations, the last 10 min of data points from the anisotropy versus time data of each reaction were averaged to give equilibrium values, then equilibrium anisotropy values versus protein concentration data were regressed with Eq. 1.2 (Sox2_HMG_-CBS and Sox2_HMG_-rG4) or 1.1 (all other interactions) to determine K_d_^app^ and n. For the displayed Sox2_HMG_ binding curves (Supplemental Figure 1b), traces were normalized to the respective maximum and minimum signals determined by regression. Mean and error (50% range) are reported in Table 1.

For initial association rate constant calculations, anisotropy versus time data for each protein concentration were fit with a smoothing spline and the initial slope of the regression was divided by the dynamic range in anisotropy for the regression to calculate apparent association rates. Apparent association rate versus protein concentration data were pruned to include only the initial linear phases, and to exclude the higher protein concentrations with incomplete curves. Then, pruned data were regressed with 0-intercept linear regression to determine k_on_^app^. Mean and error (50% range) are reported in Table 1.

For the monophasic or biphasic association fits in Supplemental Figure 3, anisotropy versus time data for each protein concentration were pruned to only include the initial data points shown in respective graphs in Supplemental Figure 3, then pruned data were regressed with Eq. 2.2 (ERα_DBD-Ext_-XBP1, Sox2_HMG_-rG4) or 2.1 (all others). Analyses were performed in R v4.3.1.

### FP-based Competitive Dissociation (FPCD)

Pre-reaction mix was prepared with 5 nM ligand in ERα (20 mM TRIS pH 7.5 at 25°C, 100 mM NaCl, 5% glycerol, 0.01% IGEPAL) or Sox2 (10 mM TRIS pH 7.5 at 25°C, 135 mM KCl, 15 mM NaCl, 0.1 mg/mL nonacetylated BSA, 4% Ficoll, 0.05% NP-40, 1 mM DTT) binding buffer, then dispensed in 32 μL volumes into the wells of a 384-well black microplate (Corning #3575). Protein was prepared at 10x the reaction concentrations of 30 nM (ERα_DBD-Ext_-dsDNA Figure 4), 100 nM (ERα_DBD-Ext_-dsDNA Figure 1), or 500 nM (ERα_DBD-Ext_-RNA and Sox2_HMG_). Competitor was prepared at 10x the reaction concentration of 10 µM (i.e., 100 µM). Microplates (ligand), protein, and competitor were then thermally equilibrated at 4°C for 30-60 min. After thermal equilibration, 4 µL of 10x protein or binding buffer (baseline control) was added to the microplate wells, then the reactions incubated at 4°C for 1 h or indicated (Figs. 4-5) shorter times. After protein-ligand incubation, 4 µL of 10x competitor or binding buffer (max signal control) was added to the microplate wells, then incubated for ≥60 min at 4°C. Fluorescence anisotropy readings were taken over the course of incubation immediately after protein addition (in <10 s intervals) with a TECAN Spark microplate reader (Ex = 481 ± 20 nm, Em = 526 ± 20 nm). Figure 1a provides helpful clarification of the methodology. Two internal controls were employed: control-1, which used buffer controls for protein addition in step-1 (Figure 1a) and competitor addition in step-2 (Figure 1a), and control-2, which used a buffer control for competitor addition in step-2 (Figure 1a). Each experiment had 2-3 reaction replicates per condition/control, and 2 independent experiments were performed per protein-polynucleotide interaction.

Anisotropy versus time data for the experimental reaction was normalized to the data for the two internal controls to give fraction bound versus time. For reactions with shorter (< 1 h) protein-ligand incubation times, an additional internal max signal control was always included where protein-ligand incubation was ≥1 hr, and this was used for normalization to calculate fraction bound. Fraction bound versus time data were regressed with Eq. 3 to determine k_fast_, k_slow_, and β_fast_. For regression, the initial fraction bound (A_max_) was constrained to the initial value calculated for the internal control with identical protein-ligand incubation that omitted competitor addition. In cases where dissociation was approximately slow monophasic, the regression was additionally constrained such that k_fast_ = 0 and β_fast_ = 0. Analyses were performed in R v4.3.1. Across experiments, percent contributions of the bi-exponential typically varied <15%.

### FP-based Jump Dilution (FPJD)

Pre-reaction mix was prepared with 50 nM ligand in binding buffer (20 mM TRIS pH 7.5 at 25°C, 100 mM NaCl, 5% glycerol, 0.01% IGEPAL), and with (experimental reaction) or without (baseline control) 50 nM protein. Wells of a 384-well black microplate (Corning #3575) were filled with 79 µL binding buffer (experimental reaction and baseline control) or 50 nM protein (max signal control), then the microplates and pre-reaction mixes were incubated at 4°C for 1 hr. After incubation, 1 µL of protein-ligand mix was diluted in the buffer-only (experimental reaction) or 50 nM protein (max signal control) microplate wells, and ligand-only mix was diluted in buffer-only microplate wells (baseline control), then the reactions incubated at 4°C for 1 h. Fluorescence anisotropy readings were taken over the course of incubation immediately after dilutions (in <10 s intervals) with a TECAN Spark microplate reader (Ex = 481 ± 20 nm, Em = 526 ± 20 nm). Two controls are employed: control-1, which uses a buffer control for protein addition, and control-2, which uses an equimolar protein control for buffer dilution. Each experiment had 1 reaction (well) per condition/control, and 3 independent experiments were performed.

Anisotropy versus time data for the experimental reaction were normalized to the data for the two internal controls to give fraction bound versus time. Fraction bound versus time data were regressed with Eq. 3 to determine k_fast_, k_slow_, and β_fast_. For regression, the initial fraction bound (A_max_) was constrained to a value of 1. Analyses were performed in R v4.3.1. Across experiments, percent contributions of the bi-exponential typically varied <15%.

### Surface Plasmon Resonance

Streptavidin-coated S-series chips were purchased commercially (Xantec #SCBS-SAD200M) and docked into a Biacore T200 SPR instrument (Cytiva). Before first-time use, all four chip flow cells (FCs) were washed (25 µL/min for 1 min) 5 times with Activation Buffer (50 mM NaOH, 1 M NaCl) to remove any unbound streptavidin, then washed (25 µL/min for 10 min) with Running Buffer (20 mM TRIS pH 7.5 at 25°C, 100 mM NaCl, 5% glycerol, 0.01% IGEPAL) to ensure surface stability. Ligand immobilization of biotin-labeled ERE (FC-2) or ΔERE (FC-4) dsDNA was performed by flowing 20 nM ligand solutions over the respective FC at 1 µL/min until a ΔRU of 300 was achieved, then priming the system with Running Buffer three times. FCs 1 and 3 were used as background controls for FC 2 and 4 experiments, respectively.

For kinetic experiments, the indicated (Supplemental Figure 4) ERα_DBD-Ext_ concentrations were flowed sequentially over the control and ligand FCs at 70 µL/min for 5 min to monitor association, followed by a 30-60 min wash phase (70 µL/min) with Running Buffer to monitor dissociation. ERα_DBD-Ext_ concentrations were tested in increasing order, and the flow cells were washed (70 µL/min for 1 min) between ERα_DBD-Ext_ concentrations 1 time with Regeneration Buffer (1 M NaCl) and 3 times with Running Buffer. Control FC signal was subtracted from experimental FC signal to generate adjusted signal versus time data, then adjusted data was exported from the instrument.

Adjusted data for each protein concentration were subjected to baseline subtraction to generate ΔRU versus time data. For initial association rate constant calculations, ΔRU versus time data were pruned to include only the association phases, then time points adjusted to start at zero time. For each protein concentration, pruned association data were used to calculate the change in ΔRU over the first 3 s of association, then this ΔΔRU was normalized to the dynamic range in ΔRU for each protein concentration to calculate apparent association rates. Apparent association rate versus protein concentration data were regressed with 0-intercept linear regression to determine k_on_^app^. For dissociation rate calculations, ΔRU versus time data were pruned to include only the dissociation phases, then time points adjusted to start at zero time. Pruned dissociation data for each protein concentration were regressed with Eq. 3 to determine k_fast_, k_slow_, and β_fast_. For regression, the initial signal (A_max_) was constrained to the final ΔRU observed at the end of the preceding SPR association phase. In cases where dissociation was approximately slow monophasic, regression was additionally constrained such that k_fast_ = 0 and β_fast_ = 0. Analyses were performed in R v4.3.1. We note that our association data have prolonged linear phases (Figure 5a), suggesting that, despite our low ligand seeding densities and high flow rates during analyte injection, mass transfer effects could be deflating the apparent association rates to some unknown degree.

### ERα Reaction Scheme Simulations

Reactions (Supplemental Figs. 6-7) were simulated and analyzed in R v4.3.1 using deSolve::ode (package::function) and the lsoda integrator (Soetaert et al. 2010). Numerical integration of the Eq. 4.1-5 system of differential equations was performed with a given integration time-step (Δt) in two phases with fixed rate constant values from Figure 6c. In phase 1 (the association phase), for initial conditions the total protein ([E_T_]) and ligand ([L_T_]) were included as free protein and ligand and all other reactants concentrations were set to zero, then the reactions were simulated for a given association time (t_on_). In phase 2, the initial conditions were set to the final reactant concentrations from phase 1 divided by a given dilution factor (d_f_), then the reactions were simulated for a given dissociation time (t_off_). Next, relative predicted anisotropy over time was calculated from reactant concentrations over time via Eq. 4.6. Equilibrium values were taken from endpoints in phase 1 simulations. Supplemental Figure 6 simulations used Δt = 25 ms, t_on_ = 1.5 h, t_off_ = 1 h, d_f_ = 10^6^, [E_T_] = 2^-12:0^ µM, [L_T_] = 5 nM, k_1_ = 10^6^ M^-1^s^-1^, k_-1_ = 1.6×10^-3^ s^-1^, k_2_ = 10^-5^ M^-1^s^-1^, k_-2_ = 3.2×10^-3^ s^-1^, k_3_ = 10^9^ M^-1^s^-1^, k_-3_ = 2×10^-2^ s^-1^, k_3b_ = 10^4^ M^-1^s^-1^, k_α_ = 2×10^7^ M^-1^s^-1^, k_β_ = 2×10^6^ M^-1^s^-1^. Supplemental Figure 7 simulations were identical, except k_-2_ = 10 s^-1^.

### Equations

For Eq. 1.1, A is the signal (anisotropy, ΔRU, fraction bound, etc.), A_min_ is the minimum signal, A_max_ is the maximum signal, E_T_ is the total protein concentration, K_d_ is the (apparent) equilibrium dissociation constant, and n is the Hill coefficient. For Eq. 1.2, L_T_ is the total ligand concentration, K_d1_ is the equilibrium dissociation constant of the first binding state, K_d2_ is the equilibrium dissociation constant of the second binding state, α is the proportion of the signal dynamic range attributable to the first binding state, and remaining parameters are as defined for Eq. 1.1.

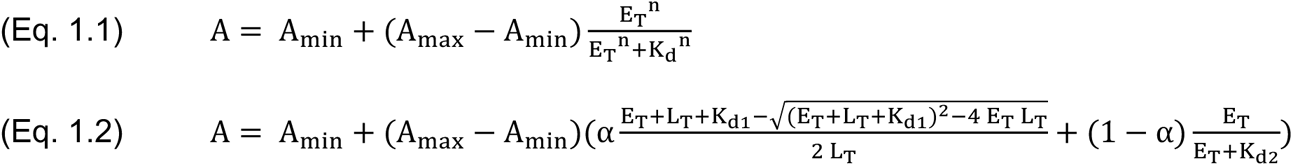

For Eq. 2.1, A_t_ is the signal (anisotropy, ΔRU, fraction bound, etc.) at a given time (t), A_min_ is the minimum signal, A_max_ is the maximum signal, k_on_ is the rate constant for the association curve, and t is time. For Eq. 2.2, k_fast_ is the rate constant for the fast phase of the association curve, k_slow_ is the rate constant for the slow phase of the association curve, β_fast_ is the proportion of the signal dynamic range attributable to the fast phase of the association curve, and remaining parameters are as defined for Eq. 2.1.

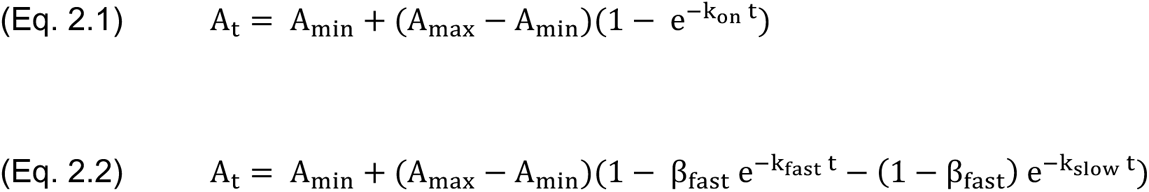

For Eq. 3, A_t_ is the signal (anisotropy, ΔRU, fraction bound, etc.) at a given time (t), A_min_ is the minimum signal, A_max_ is the maximum signal, t is time, k_fast_ is the rate constant for the fast phase of the dissociation curve, k_slow_ is the rate constant for the slow phase of the dissociation curve, and β_fast_ is the proportion of the signal dynamic range attributable to the fast phase of the dissociation curve.

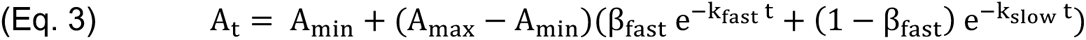

For Eq. 4, rate constants are defined in Figure 6c, E is protein, L is ligand, conjugations of these reactants are complexes, equations give rates of change for indicated reactants as a function of time (t), bracketed terms indicate concentrations, and A_rel_ is relative predicted anisotropy.

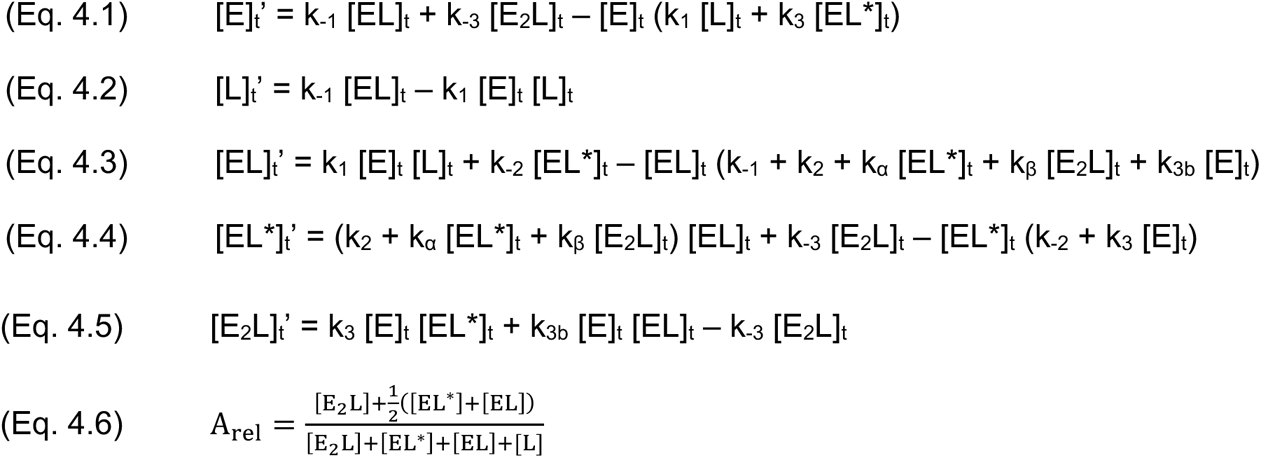

### Diagram, Reaction Scheme, and Figure Generation

All diagrams and reaction schemes were prepared on BioRender.com, tables were prepared with Word (Microsoft), graphs were prepared with R v4.3.1, and figures were assembled in PowerPoint (Microsoft).

### Data, Materials, and Software Availability

The bacterial expression plasmid for Sox2_HMG_ is available from the lab of Robert Batey (University of Colorado Boulder, Department of Biochemistry). Our bacterial expression plasmids for ERα_DBD(-Ext)_ are available upon request (contact D.S.W.). Our R script for simulating the Figure 6c reactions is available on GitHub (github.com/whemphil/ER-Sox2_Manuscript).

## ACKNOWLEDGEMENTS

W.O.H. was supported by the National Institutes of Health (F32 GM147934). D.S.W. and H.R.S. were supported by the National Institutes of Health (R01 GM120347). T.R.C. is an investigator of the Howard Hughes Medical Institute.

We thank the Batey lab (University of Colorado Boulder) for providing the Sox2_HMG_ expression plasmid and for stimulating discussion and feedback throughout these studies. We also thank Annette Erbse and the Biochemistry Shared Instruments Pool (SIP) core facility (RRID SCR_018986) for technical assistance and equipment use during these studies. We also thank the Structural Biology and Biophysics core facilities at University of Colorado Anschutz Medical campus for SPR instrumentation use and Robb Welty for technical assistance and stimulating kinetics discussions.

## COMPETING INTERESTS STATEMENT

T.R.C declares consulting status for Storm Therapeutics, Eikon Therapeutics, and SomaLogic. The other authors have no competing interests to declare.

## REFERENCES

Alluri PG, Speers C, Chinnaiyan AM. 2014. Estrogen receptor mutations and their role in breast cancer progression. Breast Cancer Res 16: 494.

Avilion AA, Nicolis SK, Pevny LH, Perez L, Vivian N, Lovell-Badge R. 2003. Multipotent cell lineages in early mouse development depend on SOX2 function. Genes Dev 17: 126–140.

Björnström L, Sjöberg M. 2005. Mechanisms of Estrogen Receptor Signaling: Convergence of Genomic and Nongenomic Actions on Target Genes. Mol Endocrinol 19: 833–842.

Chassaing N, Causse A, Vigouroux A, Delahaye A, Alessandri J-L, Boespflug-Tanguy O, Boute-Benejean O, Dollfus H, Duban-Bedu B, Gilbert-Dussardier B, et al. 2014. Molecular findings and clinical data in a cohort of 150 patients with anophthalmia/microphthalmia. Clin Genet 86: 326–334.

Chew J-L, Loh Y-H, Zhang W, Chen X, Tam W-L, Yeap L-S, Li P, Ang Y-S, Lim B, Robson P, et al. 2005. Reciprocal Transcriptional Regulation of Pou5f1 and Sox2 via the Oct4/Sox2 Complex in Embryonic Stem Cells. Mol Cell Biol 25: 6031–6046.

Crenshaw E, Leung BP, Kwok CK, Sharoni M, Olson K, Sebastian NP, Ansaloni S, Schweitzer-Stenner R, Akins MR, Bevilacqua PC, et al. 2015. Amyloid Precursor Protein Translation Is Regulated by a 3’UTR Guanine Quadruplex. PLoS ONE 10: e0143160.

De Angelis RW, Maluf NK, Yang Q, Lambert JR, Bain DL. 2015. Glucocorticoid Receptor–DNA Dissociation Kinetics Measured *in Vitro* Reveal Exchange on the Second Time Scale. Biochemistry 54: 5306–5314.

Deroo BJ, Korach KS. 2006. Estrogen receptors and human disease. J Clin Invest 116: 561–570.

Dodonova SO, Zhu F, Dienemann C, Taipale J, Cramer P. 2020. Nucleosome-bound SOX2 and SOX11 structures elucidate pioneer factor function. Nature 580: 669–672.

Fantes J, Ragge NK, Lynch S-A, McGill NI, Collin JRO, Howard-Peebles PN, Hayward C, Vivian AJ, Williamson K, van Heyningen V, et al. 2003. Mutations in SOX2 cause anophthalmia. Nat Genet 33: 462–463.

Favaro R, Valotta M, Ferri ALM, Latorre E, Mariani J, Giachino C, Lancini C, Tosetti V, Ottolenghi S, Taylor V, et al. 2009. Hippocampal development and neural stem cell maintenance require Sox2-dependent regulation of Shh. Nat Neurosci 12: 1248–1256.

Fersht A. 1985. Enzyme Structure and Mechanism. 2nd Ed. W. H. Freeman and Co., New York.

Frietze S, Farnham PJ. 2011. Transcription Factor Effector Domains. Subcell Biochem 52: 261–277.

G Hendrickson D, Kelley DR, Tenen D, Bernstein B, Rinn JL. 2016. Widespread RNA binding by chromatin-associated proteins. Genome Biol 17: 28.

Grosschedl R, Giese K, Pagel J. 1994. HMG domain proteins: architectural elements in the assembly of nucleoprotein structures. Trends Genet 10: 94–100.

Halford SE, Marko JF. 2004. How do site-specific DNA-binding proteins find their targets? Nucleic Acids Res 32: 3040–3052.

Hamilton DJ, Hein AE, Holmes ZE, Wuttke DS, Batey RT. 2022. The DNA-binding High Mobility Group Box domain of Sox family proteins directly interacts with RNA in vitro. Biochemistry 10.1021/acs.biochem.2c00218.

Han Z, Li W. 2022. Enhancer RNA: What we know and what we can achieve. Cell Prolif 55: e13202.

Helsen C, Kerkhofs S, Clinckemalie L, Spans L, Laurent M, Boonen S, Vanderschueren D, Claessens F. 2012. Structural basis for nuclear hormone receptor DNA binding. Mol Cell Endocrinol 348: 411–417.

Hemphill WO, Voong CK, Fenske R, Goodrich JA, Cech TR. 2023. Multiple RNA- and DNA-binding proteins exhibit direct transfer of polynucleotides with implications for target-site search. Proc Natl Acad Sci 120: e2220537120.

Hewitt SC, Korach KS. 2018. Estrogen Receptors: New Directions in the New Millennium. Endocr Rev 39: 664– 675.

Holmes ZE, Hamilton DJ, Hwang T, Parsonnet NV, Rinn JL, Wuttke DS, Batey RT. 2020. The Sox2 transcription factor binds RNA. Nat Commun 11: 1805.

Hou L, Srivastava Y, Jauch R. 2017. Molecular basis for the genome engagement by Sox proteins. Semin Cell Dev Biol 63: 2–12.

Hou L, Wei Y, Lin Y, Wang X, Lai Y, Yin M, Chen Y, Guo X, Wu S, Zhu Y, et al. 2020. Concurrent binding to DNA and RNA facilitates the pluripotency reprogramming activity of Sox2. Nucleic Acids Res 48: 3869–3887.

Hudson WH, Ortlund EA. 2014. The structure, function and evolution of proteins that bind DNA and RNA. Nat Rev Mol Cell Biol 15: 749–760.

Ignatieva EV, Levitsky VG, Kolchanov NA. 2015. Human Genes Encoding Transcription Factors and Chromatin-Modifying Proteins Have Low Levels of Promoter Polymorphism: A Study of 1000 Genomes Project Data. Int J Genomics 2015: 260159.

Jia M, Dahlman-Wright K, Gustafsson J-Å. 2015. Estrogen receptor alpha and beta in health and disease. Best Pract Res Clin Endocrinol Metab 29: 557–568.

Kelberman D, Rizzoti K, Avilion A, Bitner-Glindzicz M, Cianfarani S, Collins J, Chong WK, Kirk JMW, Achermann JC, Ross R, et al. 2006. Mutations within Sox2/SOX2 are associated with abnormalities in the hypothalamo-pituitary-gonadal axis in mice and humans. J Clin Invest 116: 2442–2455.

Khalil AM, Guttman M, Huarte M, Garber M, Raj A, Rivea Morales D, Thomas K, Presser A, Bernstein BE, van Oudenaarden A, et al. 2009. Many human large intergenic noncoding RNAs associate with chromatin-modifying complexes and affect gene expression. Proc Natl Acad Sci U S A 106: 11667–11672.

Kuntz MA, Shapiro DJ. 1997. Dimerizing the Estrogen Receptor DNA Binding Domain Enhances Binding to Estrogen Response Elements*. J Biol Chem 272: 27949–27956.

Lane AN, Chaires JB, Gray RD, Trent JO. 2008. Stability and kinetics of G-quadruplex structures. Nucleic Acids Res 36: 5482–5515.

Mishra K, Kanduri C. 2019. Understanding Long Noncoding RNA and Chromatin Interactions: What We Know So Far. Non-Coding RNA 5. https://www.ncbi.nlm.nih.gov/pmc/articles/PMC6958424/ (Accessed February 14, 2024).

Moosa MM, Tsoi PS, Choi K-J, Ferreon ACM, Ferreon JC. 2018. Direct Single-Molecule Observation of Sequential DNA Bending Transitions by the Sox2 HMG Box. Int J Mol Sci 19. https://www.ncbi.nlm.nih.gov/pmc/articles/PMC6321608/ (Accessed March 18, 2024).

Nair SJ, Yang L, Meluzzi D, Oh S, Yang F, Friedman MJ, Wang S, Suter T, Alshareedah I, Gamliel A, et al. 2019. Phase separation of ligand-activated enhancers licenses cooperative chromosomal enhancer assembly. Nat Struct Mol Biol 26: 193–203.

Nassa G, Giurato G, Salvati A, Gigantino V, Pecoraro G, Lamberti J, Rizzo F, Nyman TA, Tarallo R, Weisz A. 2019. The RNA-mediated estrogen receptor α interactome of hormone-dependent human breast cancer cell nuclei. Sci Data 6: 173.

Ng S-Y, Johnson R, Stanton LW. 2012. Human long non-coding RNAs promote pluripotency and neuronal differentiation by association with chromatin modifiers and transcription factors. EMBO J 31: 522–533.

Nowling TK, Johnson LR, Wiebe MS, Rizzino A. 2000. Identification of the Transactivation Domain of the Transcription Factor Sox-2 and an Associated Co-activator *. J Biol Chem 275: 3810–3818.

Oksuz O, Henninger JE, Warneford-Thomson R, Zheng MM, Erb H, Vancura A, Overholt KJ, Hawken SW, Banani SF, Lauman R, et al. 2023. Transcription factors interact with RNA to regulate genes. Mol Cell 83: 2449–2463.e13.

Parsonnet NV, Lammer NC, Holmes ZE, Batey RT, Wuttke DS. 2019. The glucocorticoid receptor DNA-binding domain recognizes RNA hairpin structures with high affinity. Nucleic Acids Res 47: 8180.

Ponglikitmongkol M, Green S, Chambon P. 1988. Genomic organization of the human oestrogen receptor gene. EMBO J 7: 3385–3388.

Rinn JL, Chang HY. 2020. Long Noncoding RNAs: Molecular Modalities to Organismal Functions. Annu Rev Biochem 89: 283–308.

Schaefer T, Lengerke C. 2020. SOX2 protein biochemistry in stemness, reprogramming, and cancer: the PI3K/AKT/SOX2 axis and beyond. Oncogene 39: 278–292.

Schwabe JWR, Chapman L, Finch JT, Rhodes D. 1993. The crystal structure of the estrogen receptor DNA-binding domain bound to DNA: How receptors discriminate between their response elements. Cell 75: 567–578.

Sisodiya SM, Ragge NK, Cavalleri GL, Hever A, Lorenz B, Schneider A, Williamson KA, Stevens JM, Free SL, Thompson PJ, et al. 2006. Role of SOX2 Mutations in Human Hippocampal Malformations and Epilepsy. Epilepsia 47: 534–542.

Skalska L, Begley V, Beltran M, Lukauskas S, Khandelwal G, Faull P, Bhamra A, Tavares M, Wellman R, Tvardovskiy A, et al. 2021. Nascent RNA antagonizes the interaction of a set of regulatory proteins with chromatin. Mol Cell 81: 2944–2959.e10.

Soetaert K, Petzoldt T, Setzer RW. 2010. Solving Differential Equations in *R*: Package **deSolve**. J Stat Softw 33. http://www.jstatsoft.org/v33/i09/ (Accessed March 15, 2024).

Spitz F, Furlong EEM. 2012. Transcription factors: from enhancer binding to developmental control. Nat Rev Genet 13: 613–626.

Steiner HR, Lammer NC, Batey RT, Wuttke DS. 2022. An Extended DNA Binding Domain of the Estrogen Receptor Alpha Directly Interacts with RNAs in Vitro. Biochemistry. 10.1021/acs.biochem.2c00536 (Accessed October 24, 2022).

Weiss MA. 2001. Floppy SOX: Mutual Induced Fit in HMG (High-Mobility Group) Box-DNA Recognition. Mol Endocrinol 15: 353–362.

Werner MS, Ruthenburg AJ. 2015. Nuclear Fractionation Reveals Thousands of Chromatin-Tethered Noncoding RNAs Adjacent to Active Genes. Cell Rep 12: 1089–1098.

Wingender E, Schoeps T, Dönitz J. 2013. TFClass: an expandable hierarchical classification of human transcription factors. Nucleic Acids Res 41: D165–170.

Wingender E, Schoeps T, Haubrock M, Dönitz J. 2015. TFClass: a classification of human transcription factors and their rodent orthologs. Nucleic Acids Res 43: D97–102.

Xu Y, Huangyang P, Wang Y, Xue L, Devericks E, Nguyen HG, Yu X, Oses-Prieto JA, Burlingame AL, Miglani S, et al. 2021. ERα is an RNA-binding protein sustaining tumor cell survival and drug resistance. Cell 184: 5215–5229.e17.

Yang F, Tanasa B, Micheletti R, Ohgi KA, Aggarwal AK, Rosenfeld MG. 2021. Shape of promoter antisense RNAs regulates ligand-induced transcription activation. Nature 595: 444–449.

Yesudhas D, Anwar MA, Panneerselvam S, Kim H, Choi S. 2017. Evaluation of Sox2 binding affinities for distinct DNA patterns using steered molecular dynamics simulation. FEBS Open Bio 7: 1750–1767.

Zhang H-M, Chen H, Liu W, Liu H, Gong J, Wang H, Guo A-Y. 2012. AnimalTFDB: a comprehensive animal transcription factor database. Nucleic Acids Res 40: D144–149.

Zhang S, Cui W. 2014. Sox2, a key factor in the regulation of pluripotency and neural differentiation. World J Stem Cells 6: 305.

